# The relationship between microbiomes and selective regimes in the sponge genus *Ircinia*

**DOI:** 10.1101/2020.08.31.275412

**Authors:** Joseph B. Kelly, David Carlson, Jun Siong Low, Tyler Rice, Robert W. Thacker

## Abstract

Sponges are often densely populated by microbes that benefit their hosts through nutrition and bioactive secondary metabolites; however, sponges must simultaneously contend with the toxicity of microbes and thwart microbial overgrowth. Despite these fundamental tenets of sponge biology, the patterns of selection in the host sponges’ genomes that underlie tolerance and control of their microbiomes are still poorly understood. To elucidate these patterns of selection, we performed a population genetic analysis on multiple species of *Ircinia* from Belize, Florida, and Panama using an *F*_*ST*_-outlier approach on transcriptome-annotated RADseq loci. As part of the analysis, we delimited species boundaries among seven growth forms of *Ircinia*. Our analyses identified balancing selection in immunity genes that have implications for the hosts’ tolerance of high densities of microbes. Additionally, our results support the hypothesis that each of the seven growth forms constitutes a distinct *Ircinia* species that is characterized by a unique microbiome. These results illuminate the evolutionary pathways that promote stable associations between host sponges and their microbiomes, and that potentially facilitate ecological divergence among *Ircinia* species.

## Introduction

Microorganisms affect nearly every aspect of macro-organismal biology. Across eukaryotes, the influences of microbiomes on hosts can be found in biological processes such as nutrition, development, and disease resistance [1–4]. These effects can be advantageous, for example, by producing essential nutrients for the hosts [5] and by enabling the exploitation of novel resources [6–8]. An important characteristic of metazoan microbiomes concerns their stability, or at the very least the prevention of their overgrowth of the host, as commensal microbes can transition into opportunistic pathogens [9]. The universal challenge of maintaining healthy associations with microbiomes is met by diverse strategies that include embargoes [10], phagocytosis [11], and physical expulsions of microbes [12]. Beneath the many mechanisms of microbiome control might lie a common process among metazoans that polices the crosstalk between hosts and their microbes: the innate immune system. This proposition is supported by observations that pathways concerned with innate immune surveillance, especially in lipopolysaccharide (LPS) sensing, are functionally conserved across distantly related host clades. LPS is a common membrane-motif found in nearly all gram-negative bacteria and has been coopted as a prototypical endotoxin that binds to toll-like receptors (TLRs), instigating an immune response. Interestingly, similarities have been documented between human and sponge LPS-induced pathways, including the likely role of toll-like receptors (TLRs) as pattern recognition receptors (PRRs) that bind LPS [13]. Furthermore, nodal signaling molecules downstream of the LPS-induced pathway in humans, serine-threonine-directed mitogen-activated protein kinases (MAPK) p38 kinases and c-*jun* N-terminal kinases/JNK [14], are also stimulated by LPS in sponges [15], a clade that is among the most distant metazoan relatives of humans. Given the homology between the components of the LPS-induced pathway, and the fact that sponges and humans share a common ancestor at the base of the metazoan phylogeny [16], these pathways might constitute extant versions of the innate immune system of ancient metazoans that facilitated inhabitation of hosts by symbiotic microbes in Earth’s early oceans, and which promote stable associations between hosts and microbes today.

Sponges stand out as a holobiont success story. The relationship between sponges and their microbes is a longstanding affair dating back to the advent of the phylum Porifera over 540 million years ago [17]. Symbiotic microbes can comprise a substantial physical portion of sponge bodies, constituting up to 40% of the total biomass in some host species [18–20]. Much like the zooxanthellae of coral, the cyanobacterial photosymbionts of sponges can supplement the host’s nutrition [4]; although in some sponges that possess high abundances of symbiotic microbes, often termed high microbial abundance or HMA sponges, cyanobacteria only constitute a portion of the microbial diversity [21]. The remaining fraction of these microbiomes can be comprised of thousands of microbial species from dozens of bacterial phyla [21] that perform fermentation [20], produce secondary metabolites [22], conduct chemoautotrophic processes such as nitrogen fixation and sulfur oxidation [23–25] and heterotrophic processes via the assimilation of dissolved organic matter (DOM) [26]. Based on measurements of nutrient transfer from microbes to their hosts [27], supplements of microbial origin to the sponges’ energy pools that the hosts use for growth [4,28], and chemical defense by the secondary metabolites [29,30], microbial symbionts can be identified as influencing the fitness of their hosts in addition to shaping their ecological identities.

The importance of microbiomes to HMA sponges is reflected in the observation that they tend to be compositionally stable and distinct relative to the microbial communities of the surrounding environment [21,31]. The microbiomes can also be divergent among host species [21] which, combined with the metabolic diversity of the microbes, supports the hypothesis that the microbiome acts as a mechanism for ecological diversification within sponges. This evolutionary model has received some of its strongest support in recent work investigating the bulk isotopic enrichment levels of sympatric sponge species [32], where both microbiome compositions and isotopic enrichment values were divergent among host taxa within geographic sites, identifying the microbiomes as being not only a mechanism for accessing novel resources but also a means for alleviating resource competition. Granted that microbiomes hold the potential to unlock access to new resources, they might also enable ecological diversification among incipient sponge species.

A suitable case study to test this hypothesis exists in *Ircinia*, a cosmopolitan sponge genus comprised of over 130 described species, many of which are densely populated by taxonomically diverse communities of symbiotic microbes [33–36]. The microbiomes of several *Ircinia* are compositionally stable [33,37] and unique among host species (Kelly & Thacker 2020 A, *in review*). These microbes also have the potential to supplement host nutrition given the high rates of primary productivity in Caribbean *Ircinia* [38] and isotopic evidence demonstrating the allocation of microbial nitrogen to the hosts [39]. In the Caribbean, *Ircinia* are characterized by divergence among the species’ microbiome compositions, a feature that might underly local adaptation (Kelly & Thacker 2020 A, *in review*). Thus, the present study sought to use Caribbean *Ircinia* to investigate how patterns of selection in the hosts’ genomes promote the residence of symbiotic microbes which could, in turn, mediate ecological divergence among closely related host species. To perform this study, we first tested whether active control of microbiomes by the hosts is evidenced throughout *Ircinia* by characterizing beta diversity among the microbial consortia of the hosts and surrounding seawater using 16S rRNA metabarcoding. Second, we tested whether this control translates to dissimilar microbiome compositions among *Ircinia* host species, several of which we delimited using 2bRAD (RADseq) data, as the possession of unique microbiomes is congruent with prior evidence of ecological diversification within sponges [32]. Finally, we identified *F*_*ST*_ outliers in the 2bRAD data and annotated them using a *de novo*-assembled and annotated transcriptome to find candidate genes that might underly divergence in microbiome control, as well as genes that could facilitate the tolerance of symbionts.

## Results

### Microbial Communities

To test whether the taxonomic compositions of *Ircinia* microbiomes are distinct relative to the community compositions of the ambient seawater microbial community, and to test whether the microbiome compositions are unique within each *Ircinia* growth form, we censused the microbial communities of seawater samples and the microbiomes of *Ircinia* spp. by sequencing the V4 region of the 16S rRNA gene. Relative abundance matrices of OTUs were constructed using mothur v1.39.5 [40] with an OTU clustering threshold of 99%. Differences among community compositions among groups (i.e. *Ircinia* and seawater) were inferred using PERMANOVA based on Bray-Curtis dissimilarity [41]. Microbiome compositions among the host *Ircinia* lineages (populations of nominal species and growth forms) were visualized using principal coordinate analysis, and dissimilarities among the microbiome compositions of these lineages were further investigated by calculating the average overlaps among the standard ellipse areas (SEAs) of the data using SIBER [42].

Sequencing of 16S rRNA amplicons generated from the host *Ircinia* and seawater samples generated 8,777,283 paired-end raw reads, of which trimmomatic removed 67.64% [43]. Of the remaining 2,839,911 reads, 2,000,318 survived downstream quality control steps including removal of homopolymers and chimeric reads in mothur v1.39.5 [40] (Table S1). 392 OTUs were identified as chloroplasts and one OTU as mitochondrial. 11042 OTUs were retrieved at the 99% clustering threshold, which SILVA v132 identified as belonging to 59 accepted and candidate prokaryotic phyla [44]. The mean final read abundance per specimen was 15127.09 +/− 7458.258 (1 SD).

The microbiomes of *Ircinia* were compositionally distinct relative to seawater microbial communities (PERMANOVA: R^2^ = 0.41, p = 1e^−04^). *Ircinia* microbiomes were 1.45x as taxonomically rich at the OTU level relative to seawater microbial communities and contained 1.56x as many source-specific OTUs relative to seawater microbial communities. The two sources only overlapped by 1043 of the 11042 OTUs, with 6115 OTUs only found in *Ircinia* and 3908 OTUs found only in seawater; however, the relative abundances of the OTUs found in both seawater and sponges were correlated with their source, where OTUs that were highly enriched in sponges were nearly absent in seawater, and vice versa (PERMANOVA: R2= 0.44, P = 1e-04, Figure 2). The shared OTUs were also the most numerically dominant in the total dataset. When considering the fraction of OTUs found in the sponges, the shared OTUs had an average relative abundance of 8.59e^−04^ +/− 4.04e^−03^(1 SD), an order of magnitude greater than the average relative abundance of sponge-specific OTUs (1.71e^−05^ +/− 1.25e^−04^ (1 SD)). The same trend held for the seawater dataset, where the shared OTUs had a mean relative abundance of 8.3e^−04^ +/− 4.42e^−03^ (1 SD), an order of magnitude greater than the average relative abundance of seawater-specific OTUs 3.35e^−05^ +/− SD 1.28e^−04^ (1 SD). 8 OTUs were identical to 16S rRNA sequences from bacteria reported to be vertically transmitted in *I. felix*, all of which fell within the intersection of the sponge and seawater datasets, 6 of which were in appreciably higher relative abundances in sponges relative to seawater (Table S4, Figure 2).

**Figure 1.**
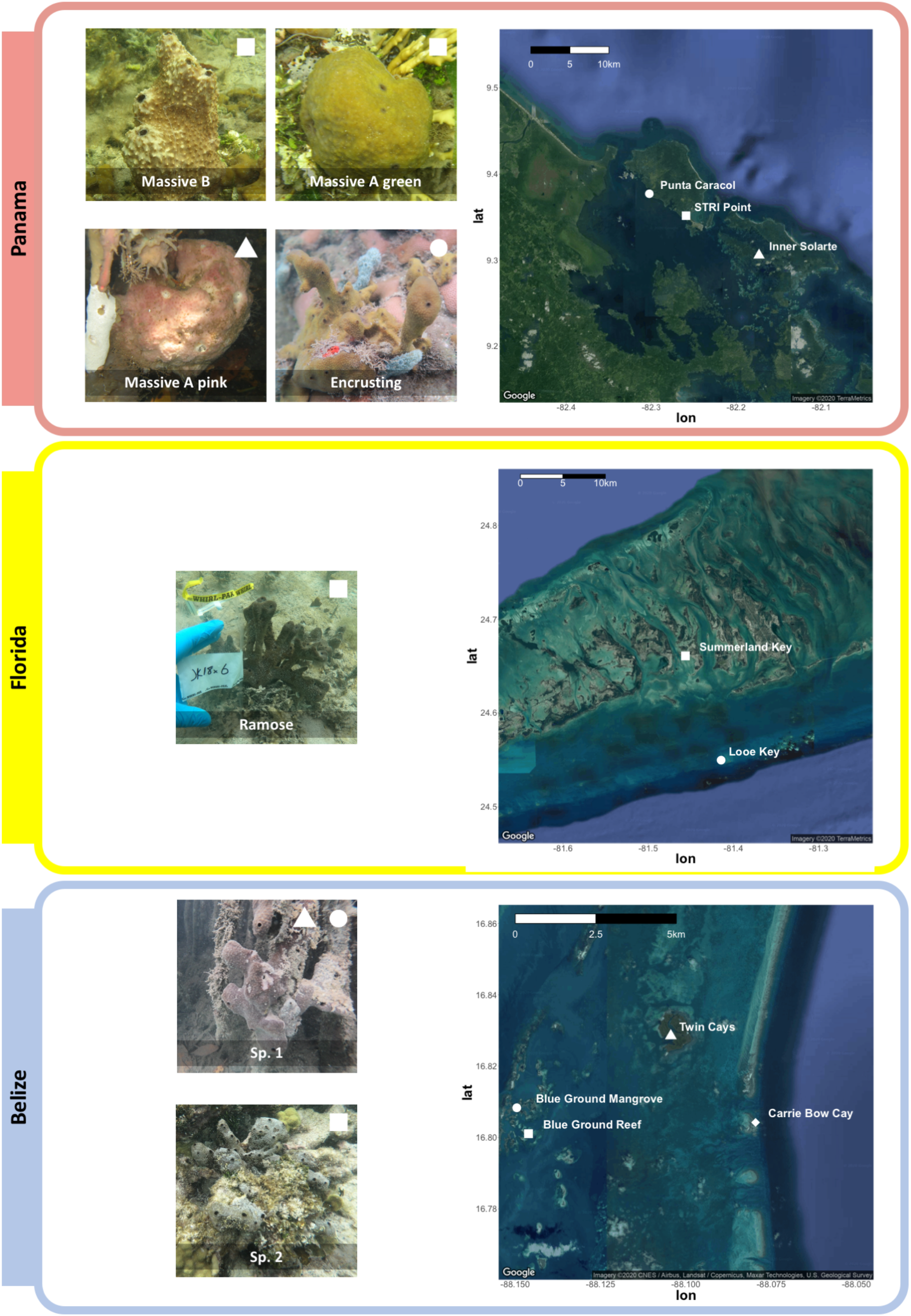
Specimens of *Ircinia* growth forms were sourced from three sites in the Caribbean. **Top:** Four Panamanian growth forms were collected from three sites: Massive A pink was collected from *Rhizophora* prop roots at Inner Solarte, a network of mangrove hammocks; Massive A green and Massive B were collected from STRI Point, a *Thalassia* seagrass-dominated habitat; and Encrusting was collected from Punta Caracol, a coral patch reef. *I. campana* and *I. strobilina* specimens were also collected from STRI Point and Punta Caracol. **Middle:** specimens of a growth form (Ramose) and *I. campana* were collected from a seagrass bed on Summerland Key, Florida; two specimens of *I. campana* were also collected from Looe Key. **Bottom:** Two Belizean growth forms were collected from three sites: Sp. 1 specimens were collected from *Rhizophora* prop roots at the Twin Cays and from mangrove hammocks adjacent to the series of Blue Ground coral patch reefs; and Sp. 2 specimens were collected from the coral reefs at Blue Ground. Specimens of *I. strobilina* were collected form the same patch reef inhabited by Sp. 2 and also from the forereef at Carrie Bow Cay, and *I. felix* specimens were collected from the Carrie Bow Cay forereef. A complete sampling overview can be found in Table S1.

**Figure 2.**
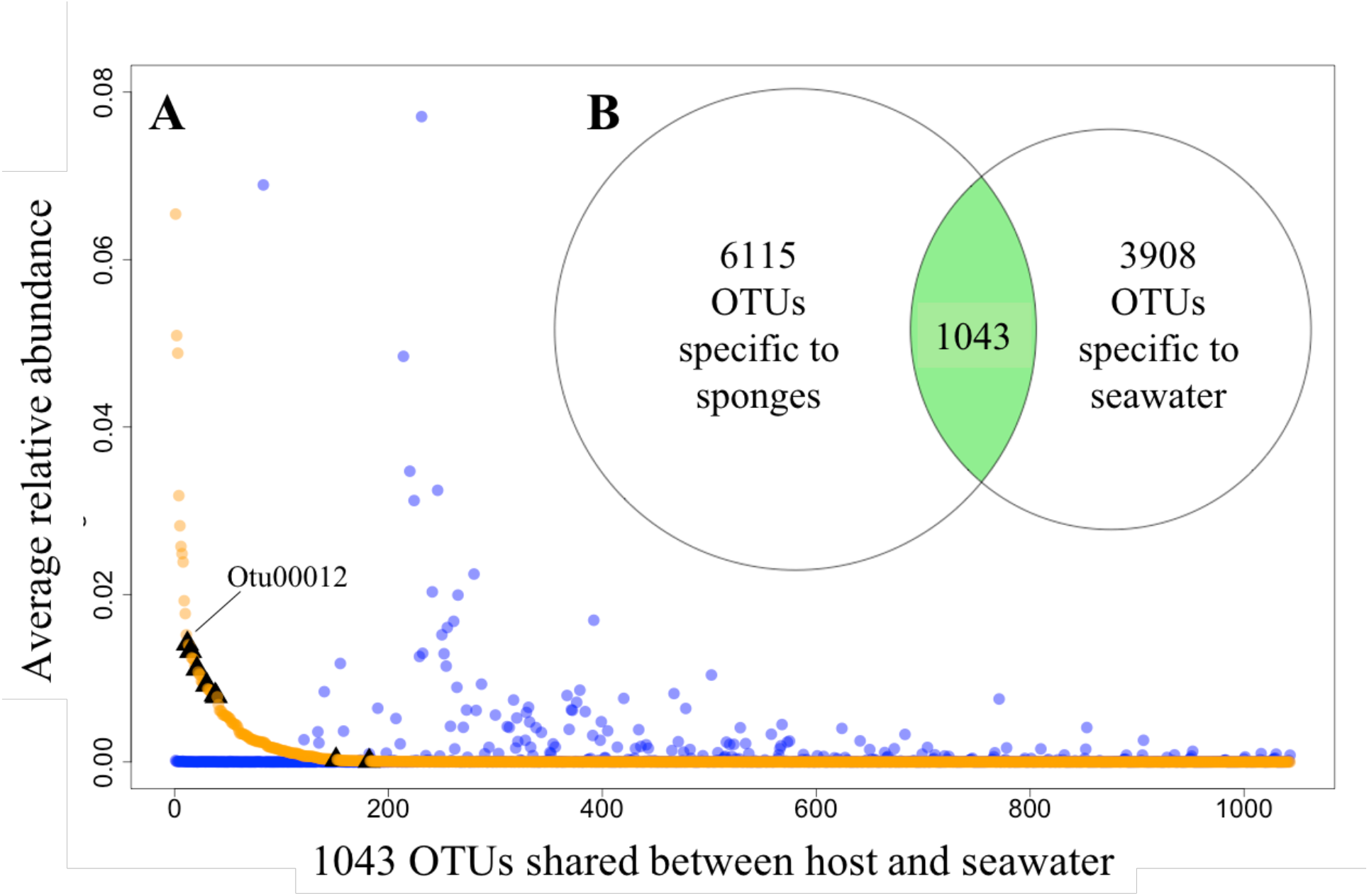
*Ircinia* possess microbiomes that are compositionally distinct relative to microbial communities of the ambient seawater. **A:** plot showing the relative abundances of the 1043 OTUs that are shared between sponges and seawater, restricted to Panama and Belize. Orange dots are relative abundances in sponges, blue dots are relative abundances in seawater. Black triangles mark OTUs that correspond to vertically transmitted bacteria in *I. felix* [58]. **B:** Venn diagram showing the number of sponge-specific OTUs, seawater-specific OTUs, and OTUs found in both sources. OTUs in the intersection of the two sources are plotted by relative abundance in A.

Each sponge lineage (growth form and population of nominal species) harbored unique microbiomes, evidenced by the significance of 66 of 67 pairwise PERMANOVAs (Table S5). Additionally, the microbiome compositions of each host lineage occupied distinct multivariate space, in which each of the standard ellipse areas (SEAs) of each host lineage had a mean overlap of 2.38 +/− 6.11 % (Figure 3). Within the nominal species *I. strobilina* and *I. campana*, geographically distant populations of conspecifics harbored significantly dissimilar microbiome compositions (Table S5). 108 OTUs were found across all 10 host species and could thus be considered ‘core’ microbiota of Caribbean *Ircinia*. One OTU that blasted to the vertically transmitted symbiont 16S sequences was found across all 10 species, Otu00012, belonging to *Constrictibacter* (Proteobacteria, Alphaproteobacteria, Puniceispirillales, Puniceispirillales incertae sedis) [45]. Of the eight putatively vertically transmitted symbionts, Otu00012 had the highest average relative abundance across all host species (Figure 2).

**Figure 3.**
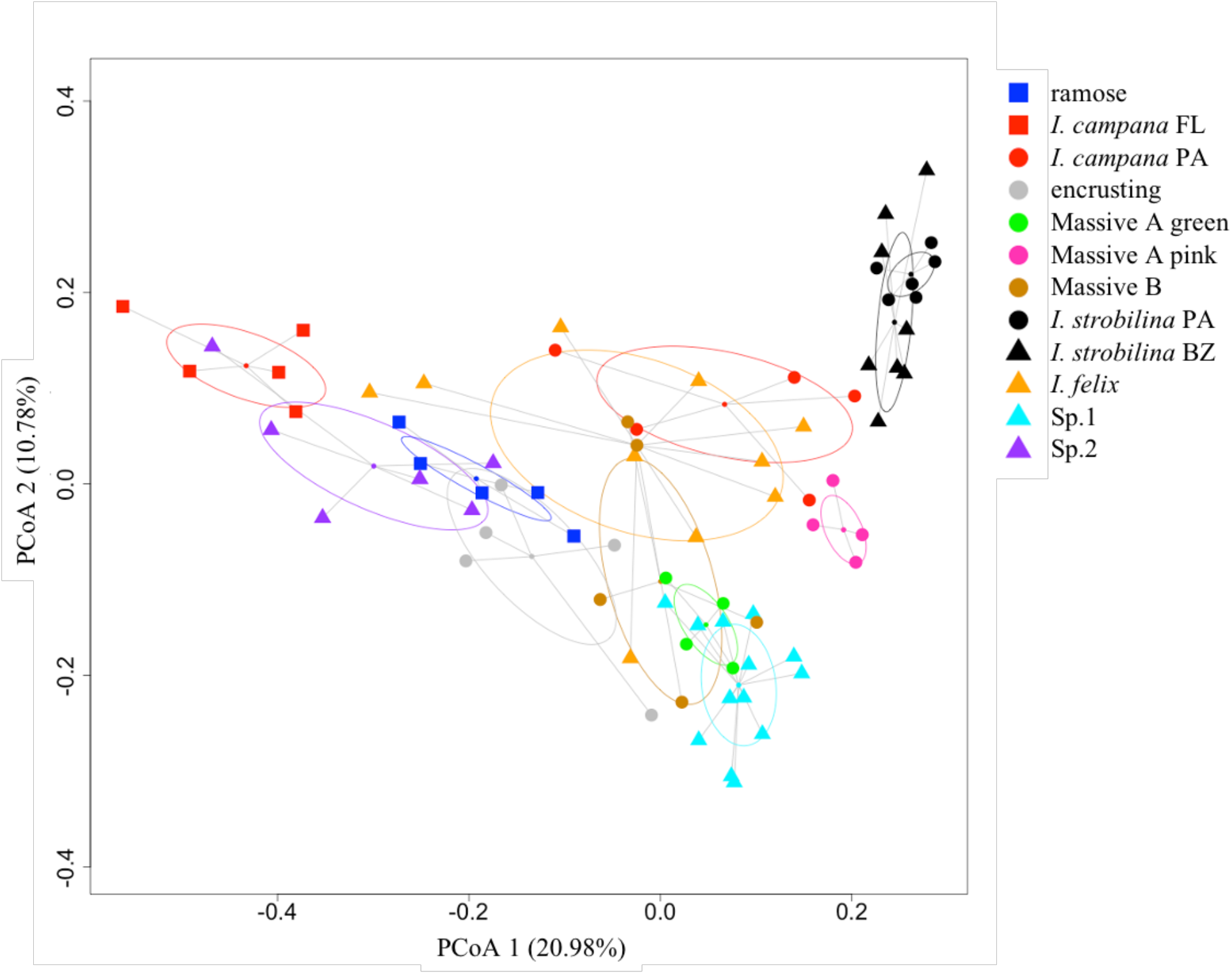
Microbiome compositions are distinct among host lineages of *Ircinia*. PCoA of microbiome compositions, normalized by relative abundance, for each host lineage. Ellipses are standard ellipse area (SEA). Squares are Floridian specimens (FL), circles are Panamanian specimens (PA), and triangles are Belizean specimens (BZ).

### F_ST_ *Outliers*

We investigated patterns of selection among the growth forms and populations of nominal species among *de novo*-assembled 2bRAD loci sourced from the host sponges using an *F*_*st*_ outlier approach. Prior to assembly and subsequent analyses, the 2bRAD data was decontaminated by removing reads that mapped to metagenome-assembled genomes (MAGs) of symbiotic prokaryotes in Caribbean *Ircinia* (Kelly et al. 2020, *in review*) using bbsplit (sourceforge.net/projects/bbmap/). A locus was identified as an outlier if it fell outside the 90% confidence interval (CI) that was estimated via fsthet [46], or if it deviated significantly from a model of neutral evolution inferred via BayeScan [47] using a 10% false discovery rate (FDR). We then annotated the outlier loci by mapping them to a *de novo*-assembled and annotated transcriptome that we produced from an *Ircinia* specimen of the Floridian ramose growth form. The transcriptome was also subjected to contaminant removal by mapping reads to *Ircinia* prokaryotic symbiont MAGs, and also using a kraken-based pipeline [48] and deconseq [49] (see Materials and Methods).

A total of 118,948,690 single-end reads corresponding to *Alf1* restriction digests were generated using the 2bRAD pipeline, with an average of 1,383,124.30 reads +/− 629,275.12 (1 SD) per specimen. After processing the reads with 2bRAD_trim_launch.pl (https://github.com/z0on/2bRAD_denovo) and cutadapt [50], a total of 103,371,805 reads remained with an average of 1,201,997.73 +/− 566,898.18 (1 SD) reads per specimen. Contamination screening against the MAGs of symbiotic prokaryotes via bbsplit removed on average 74.52% +/− 7.89% (1 SD) of reads per sample. Samples with less than 125,988.63 reads (the per-sample mean of post-decontamination reads minus 1 SD) were omitted from further analysis. 66 samples remained with an average of 360,502.20 +/− 180,813.27 (1 SD) reads per sample (Table S1). Assembly of the 2bRAD data in STACKS [51] requiring that a SNP be present in 75% of the populations and in half of the individuals per population produced 389 loci, 333 of which were variant, 50 of which were later identified as outliers by fsthet [46] and BayeScan [47].

Decontamination of the transcriptomic data via kraken [48] removed 14.88% of reads and bbsplit removed a further 0.49%. After *de novo* assembly in Trinity [52], deconseq [49] removed 3791 sequences corresponding to 1.84% of the assembled transcripts. The final assembled transcriptome had a contig N50 of 1570 bps and a metazoan-specific BUSCO completeness score of 95.6%, 191,399 transcripts, 126,897 Trinity “genes”, and a GC content of 41.58 %. 51,471 transcripts received functional annotations via HMMER, implemented in dammit [53], that met the e-value cutoff of <1e-5.

Fifty outlier loci were detected among the 333 variant 2bRAD loci; BayeScan identified 13, 10 of which were candidates for positive selection and 3 for balancing selection, and fsthet identified 43, 18 of which for candidates for positive selection and 25 for balancing selection. One locus identified as being under balancing selection and five outlier loci under positive selection were detected by both methods. 18 of these loci mapped to the transcriptome and passed the annotation criteria (Table 2). Two of the positively selected outliers mapped to genes involved in cellular mechanics (FLNC and FLNB), and one to a gene involved in maintaining DNA integrity (BLM). One of the genes under balancing selection is involved in protein degradation (S8 Family Serine Peptidase), and five in host immune and stress responses (MAP3K, Rassf1, TES, PRSS21, and Kdm5b). One of the positively selected loci mapped to a mobile genetic element (TY3B-G) as did six loci under balancing selection (three POL, gag-pol, GIY-YIG, pro-pol-dUTPase). Additionally, the two genes annotated as TES and PRSS21 also contained viral recombination domains (phage integrase family); PRSS21 also contained a reverse transcriptase domain.

**Table 1.** The species model representing each growth from and each population of nominal species as a distinct species (in bold letter face) received decisive support over competing models. Bayes factor delimitation results. Rank corresponds to relative support via Bayes factor.

| Species Model | Motivation | # Species | MLE | BF | Rank |
| --- | --- | --- | --- | --- | --- |
| **all growth forms separate species, split *I. campana* and *I. strobilina* by geography** | **Test population-level divergence.** | **12** | **-10712.77** | **2559.69** | **1** |
| split *I. campana* and *I. strobilina* by geography, combine Massive A pink and Panamanian *I. campana* | Test if Massive A pink is a Panamanian *I. campana* phenotype | 11 | -10723.72 | 2537.80 | 2 |
| all growth forms separate species, split *I. campana* by geography | Test population-level divergence. | 11 | -10743.48 | 2498.29 | 3 |
| split *I. campana* and *I. strobilina* by geography, combine Massive B and Panamanian *I. campana* | Test if Massive B is a Panamanian *I. campana* phenotype | 11 | -10743.96 | 2497.31 | 4 |
| split *I. campana* and *I. strobilina* by geography, combine Encrusting and Panamanian *I. campana* | Test if Encrusting is a Panamanian *I. campana* phenotype | 11 | -10745.95 | 2493.33 | 5 |
| split *I. campana* and *I. strobilina* by geography, combine Encrusting, Massive A pink, and Panamanian *I. campana* | Test if Encrusting and Massive A pink are Panamanian *I. campana* phenotypes | 10 | -10753.99 | 2477.27 | 6 |
| split *I. campana* and *I. strobilina* by geography, split Massive B between Panamanian *I. campana* and Encrusting (reference Kelly & Thacker 2020 A) | Test if Massive B is a mixture of Encrusting and Panamanian *I. campana* | 11 | -10767.13 | 2450.98 | 7 |
| split *I. campana* and *I. strobilina* by geography, combine Massive A pink, Massive B, Encrusting, and Panamanian *I. campana* | Test if Massive B, Encrusting, and Massive A pink are Panamanian *I. campana* phenotypes | 9 | -10779.35 | 2426.54 | 8 |
| split *I. campana* and *I. strobilina* by geography, combine sp2 and Belizean *I. strobilina* | Test if sp 2 is a Belizean *I. strobilina* phenotype | 11 | -10809.40 | 2366.44 | 9 |
| all growth forms separate species, split *I. strobilina* by geography | Test population-level divergence. | 11 | -10895.09 | 2195.06 | 10 |
| all growth forms separate species, combine *I. campana* and *I. strobilina* populations | Full species model combining populations of nominal species. | 10 | -10924.28 | 2136.68 | 11 |
| combine Massive A pink and Massive B | Both massive; shared geography. | 9 | -10940.90 | 2103.44 | 12 |
| combine Massive B and Massive A green | Shared habitat (STRI point); sympatry. | 9 | -10942.36 | 2100.53 | 13 |
| combine Massive A green and Encrusting | Sympatry | 9 | -10990.90 | 2003.45 | 14 |
| combines sp1 and Massive A pink | Shared habitat type (mangrove) | 9 | -11064.83 | 1855.58 | 15 |
| combine ramose and *I. campana* | Sympatry | 9 | -11073.93 | 1837.39 | 16 |
| combine sp2 and Encrusting | Shared habitat type (coral patch reef) | 9 | -11224.58 | 1536.08 | 17 |
| combine sp1 and sp2 | Sympatry | 9 | -11357.66 | 1269.93 | 18 |
| combine growth forms into one species | Are the growth forms different phenotypes of the same species? | 4 | -11826.39 | 332.46 | 19 |
| combine growth forms<br>with *I. felix* | Are the growth<br>forms different *I.*<br>*felix* phenotypes? | 3 | -11992.62 | - | 20 |

**Table 2.**
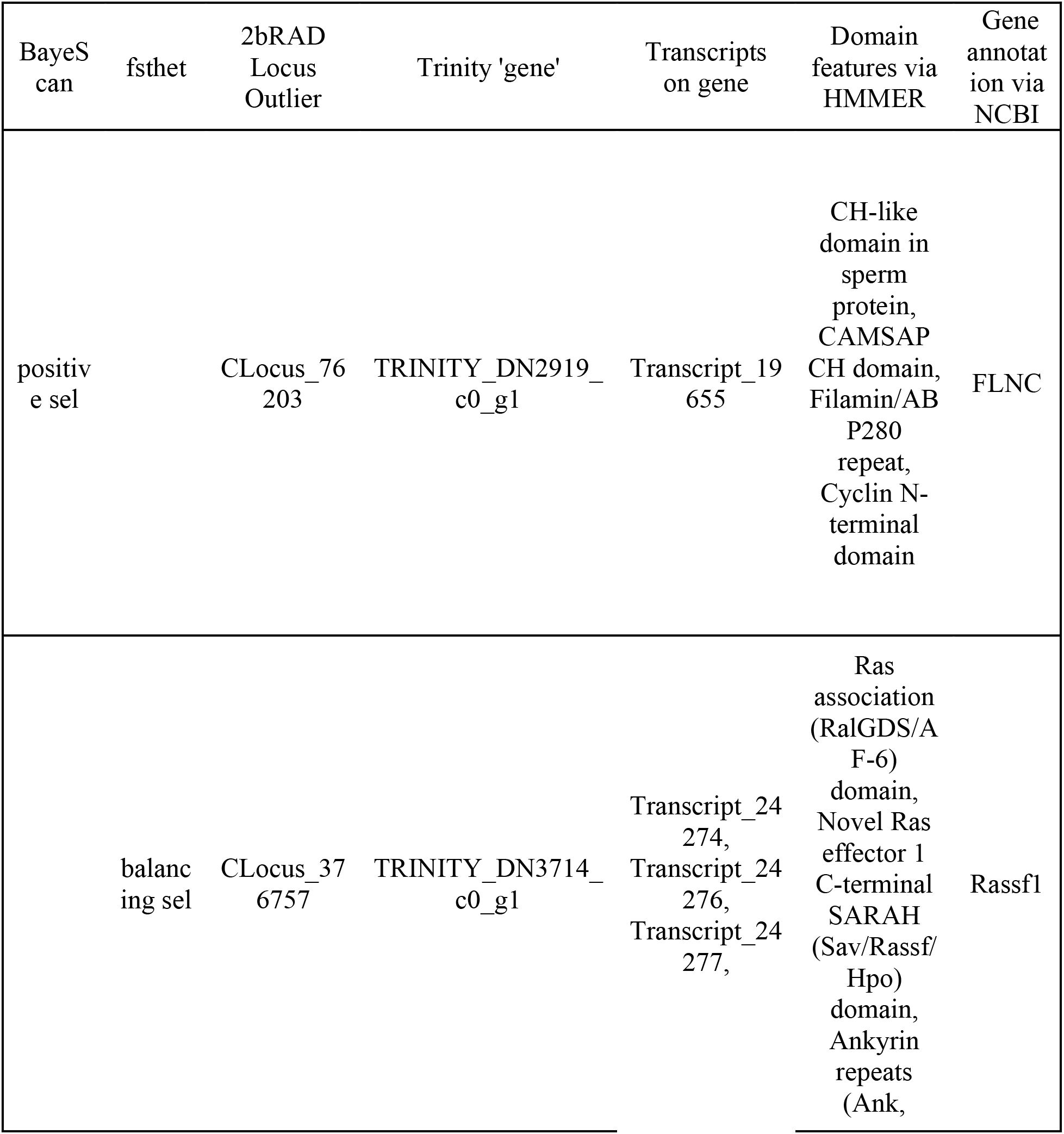
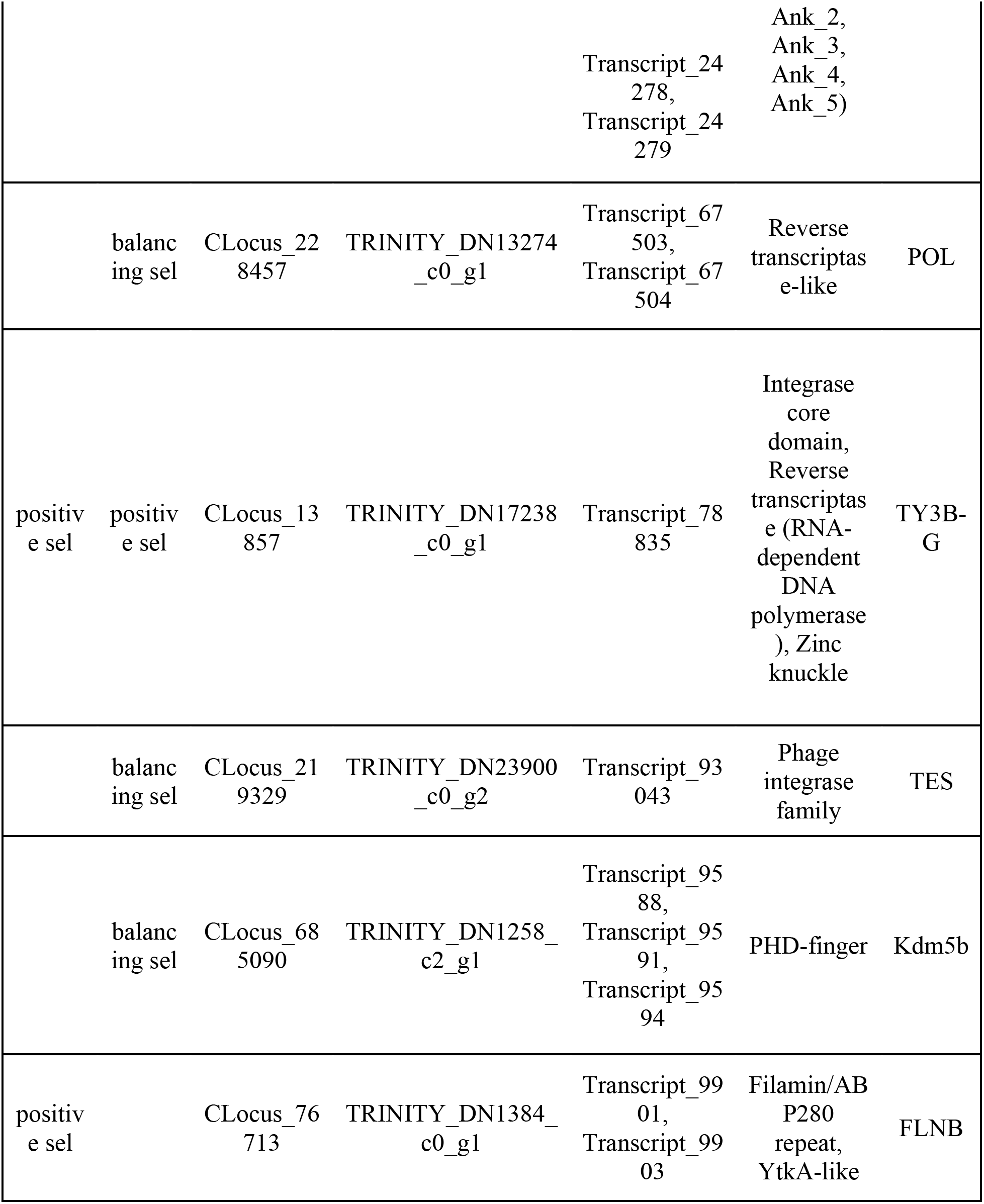
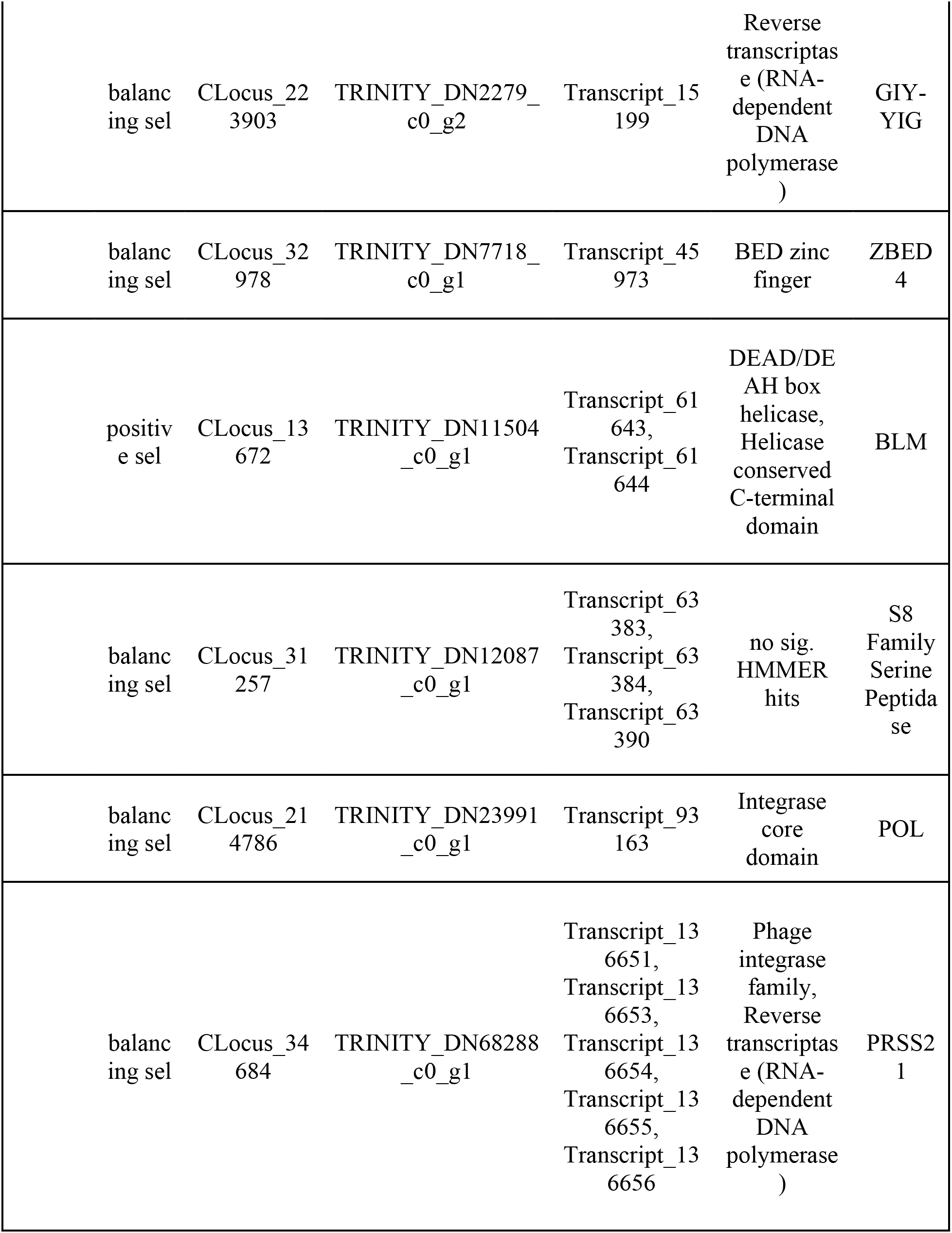
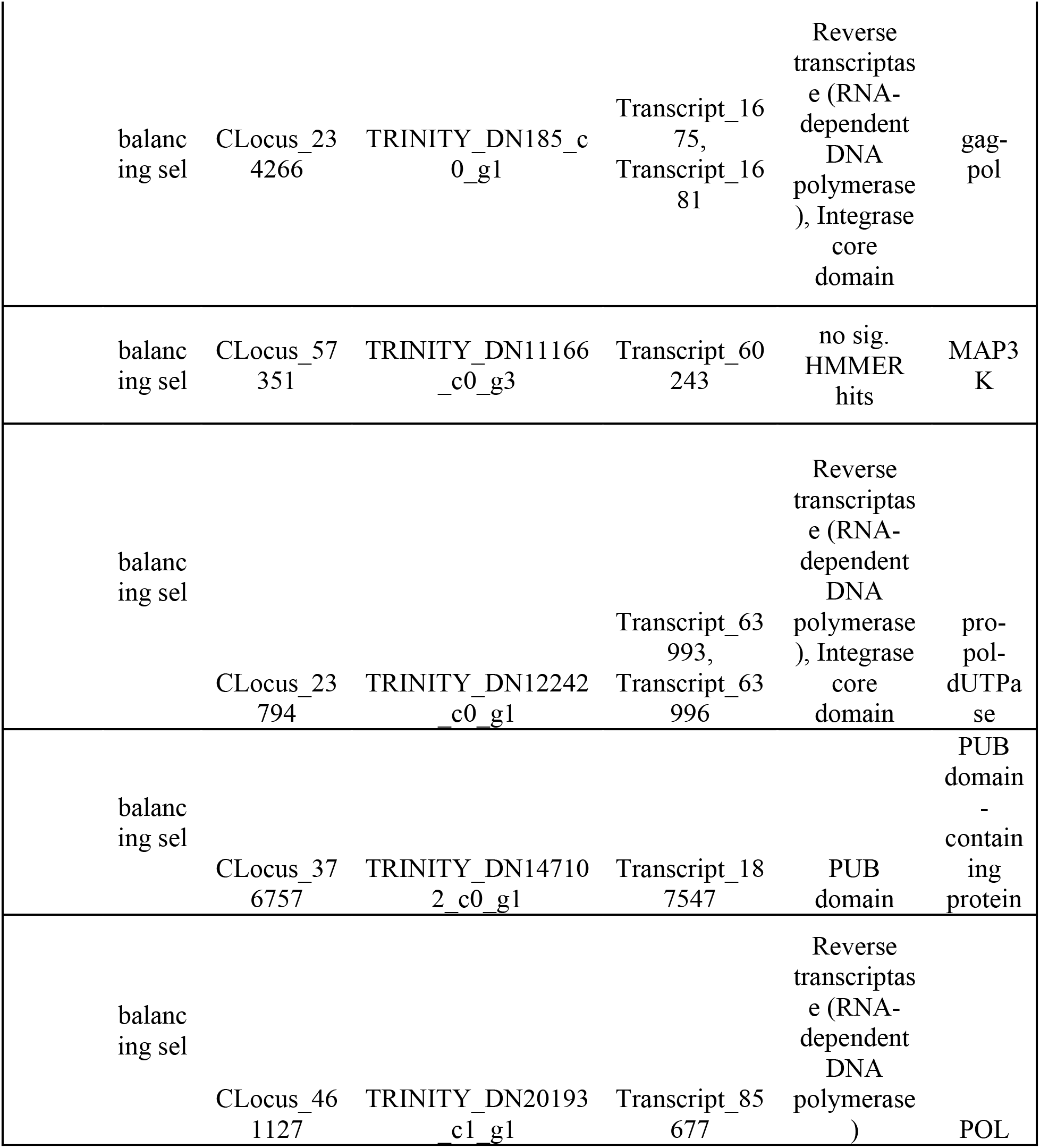
Outliers mapped to annotated transcripts. Transcripts with identical annotations are collapsed into one row. Genes involved in immunity are under balancing selection and genes involved in cellular mechanics are under positive selection in *Ircinia*. Table listing *F*_*ST*_ outliers mapped to annotated transcripts.

### Species Boundaries and Gene Flow

We delimited genetic species boundaries among *Ircinia* growth forms by performing a multispecies coalescent-based test of genetic species boundaries using BFD* [54], a pipeline that evaluates support among competing species-grouping hypotheses using Bayes factors [55]. BFD* also provides a suitable test of genetic species boundaries given our dataset as the pipeline is robust to SNP under-sampling and the sampling of relatively few individuals per species [54]. Competing species-grouping models were constructed based on whether growth forms shared a habitat type (e.g. mangrove, seagrass bed, coral reef), had similarities in the overall features of their growth habit (e.g. encrusting, massive), or were sourced from the same or nearby sites (Table 1). Additionally, we investigated patterns of hybridization among the species identified by BFD* using STRUCTURE [56]. Prior to these analyses, *F*_*st*_ outliers that were identified by either fsthet [46] or BayeScan [47] were removed to help ensure the assumption of neutrality was met [57].

BFD* lent highest support to the model representing each growth form and population of nominal species as a separate species (Table 1). The clades that were recovered via SNAPP showed a strong correlation with geography, especially with the Panamanian populations, which were monophyletic with the exception of *I. strobilina* (Figure 4). The Floridian and Panamanian *I. campana* were divided into two distinct clades that were connected by a deep node near the base of the tree; however, the *I. strobilina* populations were monophyletic.

**Figure 4.**
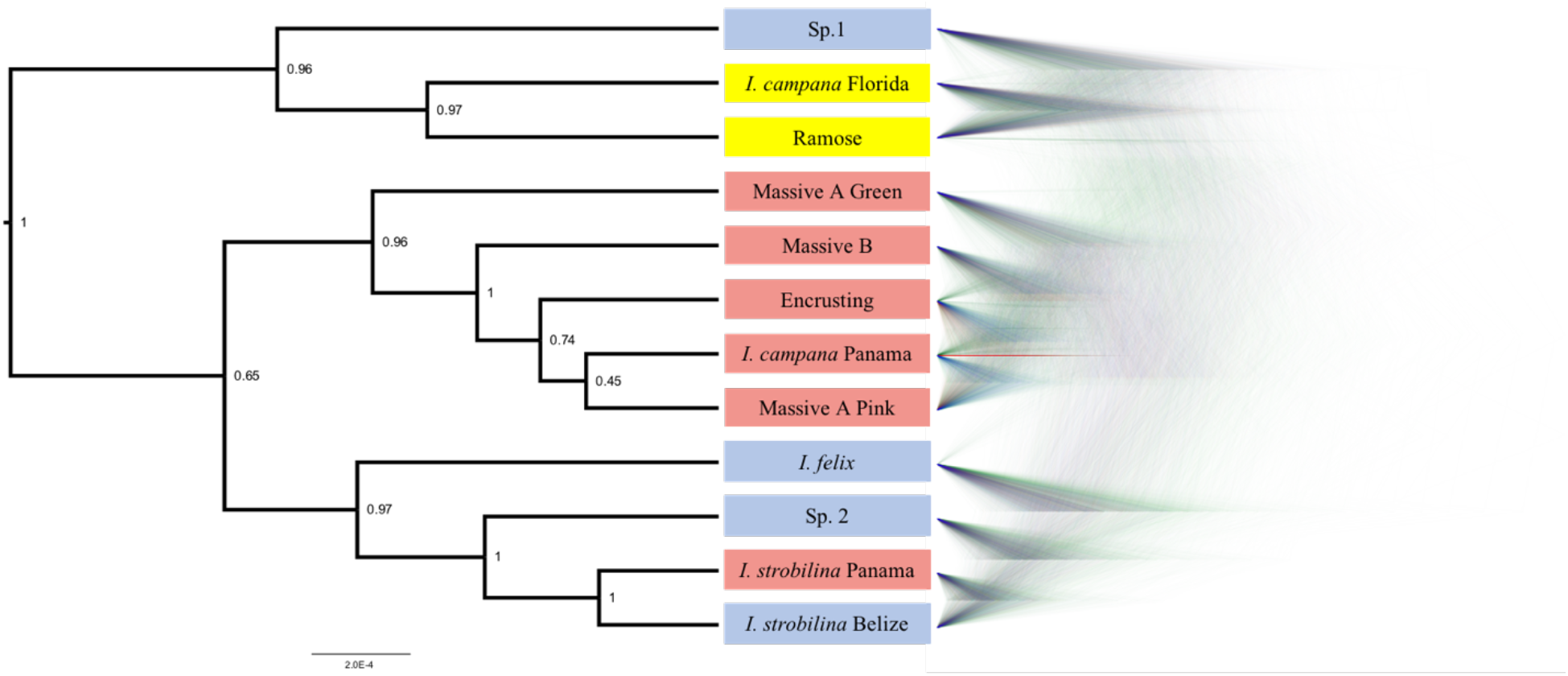
The *Ircinia* growth forms are supported as being genetically distinct species. Phylogeny produced via SNAPP for the best-supported species grouping model in BFD*. **Left:** consensus tree with posterior probabilities as node labels. **Right:** Densitree visualization of posterior tree distribution displaying most frequent topology in blue and alternative topologies in green and red. Tip labels are colored by geography: blue for Belize, yellow for Florida, and red for Panama.

The best supported number of ancestral populations was identified unambiguously as K=4, followed distantly by K=5, by the Evanno method [56] (Figure S1). Both the K=4 and K=5 STRUCTURE plots showed patterns of hybridization indicative of interbreeding among *Ircinia* species, with rates of hybridization greater within sites relative to across sites (Figure 5). In both plots, SNPs from both populations of *I. strobilina* were predominantly sourced from a shared ancestral population, whereas *I. campana* was split between two ancestral populations that coincided with geography.

**Figure 5.**
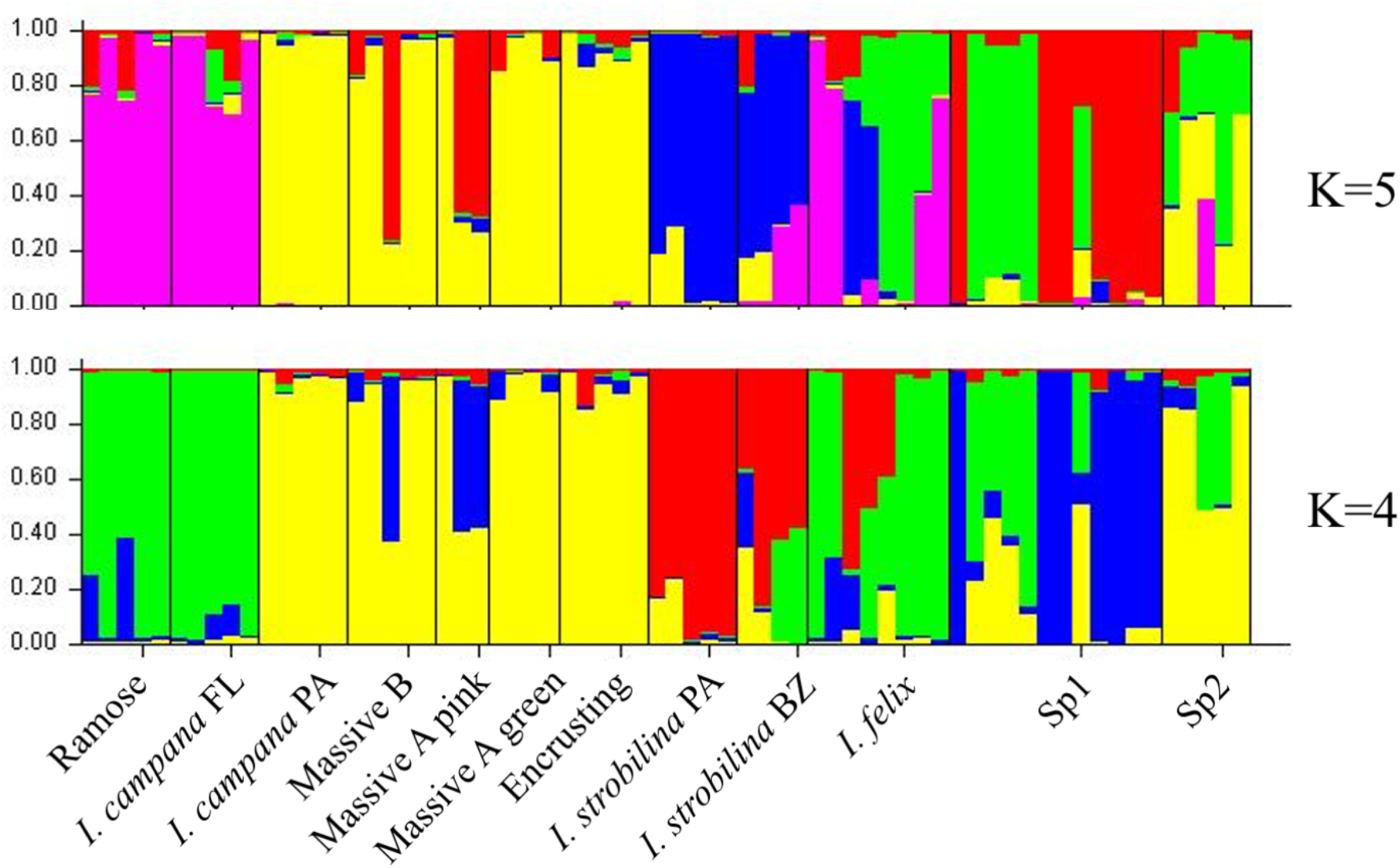
*Ircinia* experience high rates of hybridization within a site. STRUCTURE plots of SNP ancestries estimated for K=4 and K=5.

## Discussion

Our results provide the first insight into the selective forces on specific genes that might underlie microbiome control and tolerance in high-microbial abundance (HMA) sponges. In particular, balancing selection was detected in immune system genes and positive selection in genes that concern the sponge’s mechanical control of its body and DNA repair. A heritable component of microbiome control might also be evidenced in our observations of dissimilarity between host microbiomes and seawater microbial communities. Combined with previous observations of vertical transmission of microbial symbionts in sponges [58–61] and the fitness benefits to host sponges that can result from microbial farming [4,39], our study provides a model that could help explain the persistence of host-microbial relationships throughout the evolutionary history of Porifera.

### Hidden Species Richness of Caribbean Ircinia

The growth forms of Caribbean *Ircinia* appear to be genetically distinct species, a hypothesis that held despite high rates of hybridization. Two notable features stand out on the species tree for the best-supported BFD* species model. The first is a strong correlation of the grouping of taxa with geography – all four Panamanian growth forms are monophyletic with the Panamanian population of *I. campana*, and the Floridian Ramose growth form is monophyletic with the Floridian population of *I. campana*. Second, and perhaps more striking, is the polyphyly of the *I. campana* populations. Conversely, the allopatric populations of *I. strobilina* are monophyletic on the tree.

Sympatric sister lineages on the phylogeny are separated by nodes as deep or deeper than the one connecting the allopatric *I. strobilina* populations (see the pairs of sister lineages: *I. campana* Florida and Ramose and Massive A pink vs Encrusting, Figure 4). We interpret the BFD* results, the depths of these nodes, and the distinct physical characteristics of each growth form as support for the hypothesis that each growth form represents a separate species, despite high rates of hybridization. We recommend further investigation into the taxonomy of *I. campana* as our results provide evidence that this taxon likely consists of multiple species. We recommend against splitting *I. strobilina* into two species; we interpret the observed divergence among samples as population-level differences, based on comparisons of the depth of the node joining the two allopatric *I. strobilina* populations to the depths of nodes joining sympatric species pairs. Complete scientific descriptions of each of these species are provided in a separate publication (Kelly & Thacker 2020 B, *in review*).

### Microbiome Compositions of Caribbean Ircinia are Distinct from Seawater Microbial Communities and are Unique among Host Species

Consistent with prior observations (Kelly & Thacker 2020 A, *in review*), the most abundant microbes are those shared between host and seawater, whereas the OTUs specific to either sponges or seawater occur in relatively low abundances. None of these shared OTUs co-occurred in high relative abundances in both sources. Instead, sponges appear to be actively excluding some microbes that occur in high abundances in seawater and fostering high relative abundances of microbes that are only found in trace abundances in seawater. The presence of the majority of putatively vertically transmitted microbes in the set of microbes being maintained at high relative abundances in sponges supports this finding.

The microbiomes of each *Ircinia* host lineage are unique with regard to taxonomic composition, as the pairwise comparisons of the microbiome compositions of all allopatric species pairs and all but one sympatric species pairs were significantly dissimilar. Combined with the distinctiveness of the compositions of sponge microbiomes relative to the seawater microbial communities, a plausible explanation might be that a heritable mechanism in the sponges underlies microbiome assembly and potentially facilitates character displacement among sympatric species in addition to local adaptation to different habitats. Under the assumption that these microbes have an impact on the fitness of their hosts, which is a likely prospect given the high concentrations of photosynthetic pigments [62] and the presence of nitrogen metabolism [24] found in Caribbean *Ircinia*, these observations fit a model whereby microbiomes provide a conduit for ecological diversification within this genus.

The microbiomes of *Ircinia* species are also likely shaped on short-term ecological timescales to some degree; however, data describing the responses of microbiome compositions in Caribbean *Ircinia* to contrasting abiotic and biotic regimes across spatial and temporal scales are in short supply and were predominantly collected from *Ircinia* inhabiting other ocean basins. One study examining the Great Barrier Reef sponge, *I. ramosa*, found that microbiome compositions were stable when exposed to different salinity regimes [63]. Additionally, previous studies focusing on Mediterranean *Ircinia* spp. discovered that microbiome demographics are stable within a host species across seasons and throughout the range of their species distributions [33,37]. Recently, an analogous study was performed in the Caribbean that investigated patterns of beta diversity among the microbiomes of *I. campana* populations [35]. This study discovered a distance-decay relationship among the microbiome compositions of the populations [35], which is consistent with the significantly dissimilar microbiome compositions of the *I. campana* populations in the current study. However, given the topology of our species tree and the BFD* results, it may be the case that more than one *Ircinia* species were investigated by the aforementioned study, again confounding interpretations of the relative contributions of species-level ecological divergence and ecological factors that scale with latitude. To better understand how intersecting evolutionary and ecological forces dictate microbiome assembly in Caribbean *Ircinia*, we advocate for more studies performing long-term monitoring of microbiome compositions in Caribbean *Ircinia* and reciprocal transplant experiments that could help elucidate how *Ircinia* are able to use their microbiomes to exploit unique resource pools in different habitats.

### The Role of Selection in Ircinia Evolution

The demands of living in a microbial world must be met by competent host defenses. Immune system genes are routinely subjects of balancing selection given the positive effect allelic diversity has in guarding against novel cellular assaults and in detecting a broader suite of foreign epitopes [64], and have been previously detected as being under balancing selection in sponges [65]. In our dataset, we detected multiple outliers under balancing selection involved in immune responses. The first two, MAP3K and Rassf1, are both expressed in the cytosol and are components of the Ras-Raf-MEK-ERK (MAPK/ERK) LPS-induced pathway that communicates molecular signals of a bacterial infection to drive the transcriptional changes necessary for an immune response [66]. This signaling pathway also plays a role in cell-cycle mediation and in the induction of apoptosis, and is thought to be tumor-suppressing given the silencing of Rassf1 and somatic mutation of MAP3K in many cancers [67–70]. Two other oncogenes under balancing selection are TES (testisin) and PRSS21 (testisin precursor). Both are localized in the cell membrane. The former is a focal adhesion protein that controls cell proliferation [71], the latter, a serine protease that is putatively involved in the regulation of proteolysis during germ line development [72]. Our analysis detected both positive and balancing selection in biological processes involved in maintaining DNA integrity. Balancing selection was detected in Kdm5b, a histone demethylase expressed in the nucleus that signals double-stranded breaks in DNA [73]. Balancing selection was also detected in a member of the GIY-YIG nuclease family, which includes members that prevent the incorporation of exogenous DNA and are thought to preserve DNA integrity in basal metazoans [74,75]. Positive selection was detected in the BLM protein, which helps ensure accurate recombination during double-strand DNA break repair [76].

The concentration of bioactive compounds in *Ircinia* can be substantial, and some that have been identified as being produced by the symbiotic microbes are cytotoxic [77–79]. Many prokaryotic symbionts present in the adult sponges have also been observed in *Ircinia* larvae, where the toxic effects of the secondary metabolites are likely pronounced [58]. Thus, balancing selection in immune system genes could be having a fitness effect in multiple cellular compartments (cell membrane, cytosol, and nucleus) and over the lifespan of the host. Additionally, the patterns of selection present in genes that promote DNA integrity, including oncogenes, could result in adaptations to the molecular apparatuses that prevent mutagenesis of host DNA in an environment that contains abundant foreign DNA and secondary metabolites.

Both balancing and positive selection were detected in viral recombination genes, providing the second account of selection in genes of a viral origin in sponge hosts [65]. Additionally, a recent study investigating the roles of phages in sponge symbionts discovered that bacteriophages of four Mediterranean sponge species contain ankyrin repeats, which likely aid the symbiont in evasion of the host’s immune system, and other genes that supplement the host bacterium’s metabolism [80]. Since similar viral loci are under selection in the host, it may be the case that viruses are also transferring adaptive gene content to the sponges. Another biologically plausible scenario is that these genes are erroneously being detected as under selection, in that if they are in close proximity to other genes that are the true targets of selection, then our analysis could be detecting them as a result of a hitchhiking effect. However, this scenario does not preclude the possibility that the viral loci might still somehow be involved in the mobility of the target of selection. Further research that can provide information on the physical relationship among these loci (i.e. using a reference genome) is required to better vet the potential for viruses to introduce adaptive genes to host sponges.

Two of the loci under positive selection (FLNC and FLNB) are involved in cellular mechanics, including the development and functioning of muscles in other metazoans [81–83]; additionally, the copy of FLNC in our dataset contains a CH-like domain that is present in sperm flagella (Table 2). These genes might be involved in contractions of the canal system and the mechanics of the flagellar beating of the choanocytes, which regulate the flow of water throughout the sponge [84]. Some host organisms, such as legumes, control their microbiome by manipulating the microenvironment, which can deprive root nodules that are overgrown with cheater strains of rhizobia of oxygen [10]. Sponges might control their microbiomes analogously, as they are able to control which portions of their aquiferous canal system receive irrigation, resulting in a heterogenous distribution of oxygen that could impact physiologies of bacterial symbionts and thus change the microbiome composition [85]. Given that the growth forms are specific with regard to habitat preference, the divergence among these genes could further be compounded by the different hydrodynamic environments of coral reefs, seagrass beds, and mangroves [86].

## Conclusion

Ecological divergence, as facilitated by the microbiomes of Caribbean *Ircinia*, could be enabled in part by the patterns of selection we detected in the genomes of the hosts, which include balancing selection at immunity genes and positive selection in genes involved in cellular mechanics and the maintenance of DNA integrity. Of special interest are the immunity genes, as the innate immune system of sponges might play a central role governing host-microbial crosstalk and maintaining a healthy microbial homeostasis [87]. Immunity pathways involving the mitogen-activated protein kinases (MAPKs) p38 protein kinase and c-*jun* N-terminal kinases/JNK are induced by LPS in the model sponge species *Suberites domuncula* [15]. One of the pathways detected as being under balancing selection here, Ras-Raf-MEK-ERK (MAPK/ERK), is stimulated by LPS in human cell lines and triggers downstream immune responses from the host [14]. Given the conservation of the actions of the other two MAPK pathways, a similar biological function could perhaps be performed by MAPK/ERK in *Ircinia*. Future work should test whether this pathway is inducible by microbes or microbial metabolites and investigate the implications of its role in the cell cycle for tolerance of the microbiome. By identifying the products of genes in this pathway and of other genes that we detected as being under selection, such research will further illuminate how sponges coexist with their microbiomes and how selection in the host genome might drive microbially mediated ecological diversification.

## Materials and Methods

### Specimen collections and next generation sequencing library preparation

Specimens of *Ircinia* representing seven growth forms were collected from three sites: Bocas del Toro, Panama (July 2016), the Florida Keys (July 2018), and the Mesoamerican Barrier Reef (August 2018) (Figure 1, Table S1). In Panama, individuals of the growth form Massive A pink were collected form mangrove prop roots of *Rhizophora* at Inner Solarte; individuals of two growth forms, Massive A green and Massive B, were collected from the seagrass-dominated (*Thalassia*) habitat of STRI point; and individuals of the Encrusting growth form were collected from patch reefs of Punta Caracol. In Florida, specimens of *I. campana* and a growth form with a branching body morphology (Ramose) were collected from a 150 m-long seagrass bed that begins 50 m immediately to the west of MOTE Marine Laboratory and Aquarium’s Elizabeth Moore International Center for Coral Reef Research & Restoration; *I. campana* specimens were also collected from the coral reef at Looe Key. An additional specimen of the Ramose growth form was snap frozen in an EtOH – dry ice bath and stored at −80C for subsequent RNA extraction. In Belize, specimens of *I. strobilina* and *I. felix* were collected from the forereef on the western (seaward) slope of Carrie Bow Cay. Specimens were collected of a sixth growth form with an irregularly massive body morphology (Sp. 1) from the *Rhizophora* prop roots of the Twin Cays and Blue Ground mangrove hammocks, and of a seventh growth form with an encrusting body morphology (Sp. 2) from the Blue Ground coral patch reef. Specimens of *I. strobilina* were also collected from the Blue Ground patch reef alongside Sp. 2. All growth forms are specific with regard to habitat preference and inhabit either coral patch reefs, seagrass beds, or mangrove prop roots, and were collected in close proximity to each other within a site; all Panamanian collections were made within a 10.7-km radius, the Floridan specimens shared a habitat, and the Belizean sampling locations fall within a 7.5-km radius.

Sponge specimens were fixed in 90% EtOH, which was replaced at the 24-hour and 48-hour marks. 0.5L seawater specimens were taken immediately adjacent to the sampled *Ircinia* in Panama and transported in opaque brown Nalgene bottles, subsequently concentrated via vacuum filtration through 0.2-μm Whatman filter papers, and then stored in RNA later. DNA extractions were made from the outermost 2mm layer of the sponge tissues using DNeasy PowerSoil Kit (Qiagen) and from the interior of the sponge tissue (<2mm from the exterior pinacoderm) and the seawater filter papers using the Wizard Genomic DNA Purification Kit (Promega).

To census the taxonomic microbial community compositions of sponges and seawater, we amplified the V4 region of the 16S rRNA (ribosomal subunit) from the DNeasy PowerSoil Kit DNA isolations using the primers 515f (5’ GTG YCA GCM GCC GCG GTA A 3’) and 806rB (5’ GGA CTA CNV GGG TWT CTA AT 3’) following the Earth Microbiome Project 16S protocol (http://press.igsb.anl.gov/earthmicrobiome/protocols-and-standards/16s/). PCR reactions were conducted in 50 uL volumes with the following recipe: 25 uL of 2x HotStarTaq Master Mix, 1 uL of each primer at 10 uM concentration, 22 uL H_2_0, and 1 uL DNA template. Thermocycler conditions used an initial denaturing step of 95°C for 5 minutes followed by 35 of the following cycles: 94°C for 45 seconds, 50°C for 1 minute, and 72°C for 1.5 minutes; and was completed with a final elongation step of 72°C for 10 minutes.

To generate genome-wide SNPs data for the host sponges, we constructed a 2bRAD (RADseq) library from the sponge DNA isolations produced using the Genomic DNA Purification Kit following the workflow of Wang et al. 2012 [88], whereby all *Alf1* restriction sites were targeted for amplification with the primers 5ILL-NN (5’ CTA CAC GAC GCT CTT CCG ATC TNN 3’) and 3ILL-NN (5’ CAG ACG TGT GCT CTT CCG ATC TNN 3’) [88]. The 16S and 2bRAD libraries were multiplexed using 12-basepair Golay barcodes and pooled within each amplicon type following dsDNA quantification on a Qubit 3.0.

RNA was extracted from the frozen Ramose specimen by incubating a homogenized tissue fragment in Trizol and processing the resultant aqueous phase through the QIAGEN RNAeasy kit following the manufacturer’s instructions. The RNA extraction was sent to the Yale Center for Genome Analysis (YCGA) for library preparation via poly-A pulldown and sequencing on a NovaSeq6000. The 16S rRNA library was sequenced on a MiSeq in the lab of Dr. Noah Palm at Yale University using a V2 2×250 bp chemistry kit, and the 2bRAD library was sequenced on a NovaSeq 6000 at YCGA.

### Read Preprocessing and Data Hygiene

Initial quality filtering was performed on the 16S rRNA reads using the paired-end function in Trimmomatic v0.36 with the settings TRAILING:30 SLIDINGWINDOW:5:20 MINLEN:100 [43]. The forward 2bRAD reads were trimmed to the 36-bp restriction fragments using the script 2bRAD_trim_launch.pl (https://github.com/z0on/2bRAD_GATK), and quality filtered using cutadapt with the settings -q 15,15 -m 36 [50]. Decontamination of the 2bRAD dataset (i.e. removal of reads of prokaryotic origin) was performed using bbsplit.sh (https://sourceforge.net/projects/bbmap) with default mapping parameters against a reference database of 356 metagenome-assembled genomes (MAGs) sourced from the same 12 *Ircinia* host lineages studied here (Kelly et al. 2020, *in review*). RNAseq reads were filtered and trimmed prior to assembly using the paired-end function of fastp v0.19.6 with the options --poly_g_min_len 10 -x --poly_x_min_len 10 [89]. A first round of contaminant removal was performed on the RNAseq data using kraken v1.1.1 with the parameter setting --confidence 0.05 [48]. To mitigate contamination of both eukaryotic and prokaryotic commensals, we supplemented the default kraken databases with custom databases built from the aforementioned dereplicated MAG dataset of *Ircinia* and a set of publicly available genomes downloaded from NCBI, which is predominantly crustacean and annelid as these taxa comprised the majority of eukaryotic commensals in the ramose growth form (Table S2). A second round of contaminant removal was performed using bbsplit.sh, implemented with a modification to default mapping parameters of maxindel=200000 and a reference database of the dereplicated *Ircinia* MAG dataset.

### 16S rRNA Analysis

16S rRNA reads passing initial quality filters were assembled into contigs, demultiplexed, and aligned to the V4 region of the SILVA v132 SSU reference sequence database in mothur v1.39.5 [40]. Following removal of chimeric sequences, OTUs were clustered at the 99% threshold using distance-based greedy clustering implemented in VSEARCH [90]. OTUs represented by only one or two reads were omitted from the final dataset to mitigate read error and contamination. Additionally, OTUs that were identified by SILVA as being from mitochondria, chloroplasts, or Eukaryotes were removed from the dataset prior to downstream analyses. To infer which OTUs might be vertically transmitted, we used BLASTn to match the representative 16S rRNA sequences of our OTUs to a database of 16SrRNA sequences from bacteria that are putatively vertically transmitted in *I. felix* [58], downloaded from NCBI (Table S3). OTUs were identified as being putatively vertically transmitted if they had 100% sequence identity over the entire length of the query.

Beta diversity among the host species and differences in microbial community compositions between sponges and water were inferred using Permutational Analysis of Variance (PERMANOVA) based on Bray-Curtis dissimilarity [41,91]. Sponge microbiome compositions were visualized in multivariate space using a PCoA, and the overlap among the standard ellipse areas of each host species’ microbiome composition was calculated using SIBER [42]. The number of unique and shared OTUs between seawater and sponge microbial communities were plotted with the eulerr package (https://github.com/jolars/eulerr). Since seawater microbial communities were not sampled from Florida, analyses comparing sponge and seawater microbial communities are restricted to Panamanian and Belizean samples.

### Transcriptome Assembly, Post-Assembly Decontamination, and Functional Annotation

Reads passing the kraken and bbsplit.sh filtering steps were *de novo* assembled using Trinity v2.8.5 [52]. After assembly, a third round of contamination removal was performed using deconseq v0.4.3 with default parameters and the publicly available mouse, human, bacterial, archaeal, and viral deconseq reference databases [92]. Functional annotations were then made for the assembly using the annotate function of dammit v1.0rc2 (https://github.com/dib-lab/dammit) including the UniRef90 annotations (--full) and the metazoan lineage-specific BUSCO group [53].

### De novo 2bRAD assembly

Following parameter optimization via the guidelines of Paris et al. [93], we assembled the 2bRAD reads into loci *de novo* in Stacks v2.41 with the settings -m 3 -M 3 -n 4 [51]. Additionally, we filtered the data to require that a SNP be present in 75% of the populations (-p 9) and half of the individuals of a population (-r 0.50). For downstream population genetic analyses, we used only the first variant site from each 2bRAD locus to satisfy the assumption of independence among our SNPs.

### *F_ST_* Outlier Detection and Annotation

Using two methods, we detected *F*_*ST*_ outliers by treating each growth form and allopatric population of nominal species as a separate population. First, via BayeScan v2.1, a Bayesian software program that employs reversible-jump Monte Carlo Markov chains to estimate posterior distributions of *F*_*ST*_ values for loci under two alternative models, one with selection and another under neutral evolution [47]. Loci with posterior *F*_*ST*_ values that deviate from expectations under the neutral model are identified as outliers. BayeScan was run for 5000 iterations with a thinning interval of 10 and a burn-in of 50000. Outliers were identified from the output files and plotted using the R function plot_bayescan with a false discovery rate (FDR) of 0.10. Second, we identified outliers using the R package fsthet, an implementation of the *F*_*ST*_-heterozygosity approach of Beaumont & Nichols (1996) that identifies outlier loci against smoothed quantiles generated from the empirical SNP dataset [46]. Using this approach, a given locus was identified as an outlier if it fell outside the 90% confidence interval (CI) of its heterozygosity bin.

To investigate the biological implications of selection in our dataset, we performed a two-step functional annotation. First, we mapped 2bRAD loci containing outliers by querying them against the assembled and annotated transcriptome using BLASTn with an e-value cutoff of 1e-9. Functional annotations of transcripts containing outlier loci were deemed reliable if the HMMER hits had e-values below 1e-5. Second, we queried outliers via BLASTx to the to the NCBI non-redundant (nr) protein sequences database.

### Species Delimitation, Species Tree Estimation, and Hybridization

SNPs that were identified as *F*_*ST*_ outliers were removed from the SNP matrix prior to downstream population genetic analyses. Species delimitation was performed using Bayes factor delimitation with genomic data (BFD*) [54]. 13 competing species grouping models were constructed based on plausible biological scenarios (Table 1). Each model was assigned an alpha = 1 and beta = 130 for the expected divergence prior θ and a prior distribution of gamma (2,200) for the birth rate prior λ (https://www.beast2.org/bfd/). Marginal likelihoods were then estimated for each competing model via path sampling analysis [95], which was run for 50,000 generations with 28 path steps and a pre-burn-in of 25,000 generations. Bayes Factors were calculated and compared following Kass & Raftery [55]. A species tree was estimated in SNAPP v1.3.0 for the best-supported species grouping model using four MCMC chains, each with a length of 1 million generations and a burn-in of 25% (totaling 3 million generations post-burn-in) [96]. Likelihood estimates and trees were logged every 500 generations for the SNAPP species tree and the path sampling analyses. Hybridization was inferred among the species identified by BFD* using STRUCTURE by setting the number of ancestral populatiions (*K*) to range from 3 to 12 and performing 10 runs for each *K* using an MCMC length of 200,000 and a burn-in of 50,000 generations [56]. *ΔK* was calculated to estimate the number of ancestral populations using the Evanno method implemented in Structure Harvester v0.6.94 [56,97].

## Data accessibility statement

The quality-controlled 16S rRNA, 2bRAD, and transcriptomic sequence data are deposited under the GenBank accession numbers XXX, YYY, ZZZ, respectively.

## Acknowledgements

We thank Dr. Jackie L. Collier and Dr. Liliana Davalos-Alvarez for their comments on the manuscript; the staffs of the Smithsonian Tropical Research Institute’s Bocas Research Station, the Smithsonian’s Carrie Bow Cay Field Station, and the MOTE Marine Laboratory and Aquarium’s Elizabeth Moore International Center for Coral Reef Research & Restoration for their support on logistical aspects of the field work; and Barrett Brooks and Dr. Karen Koltes for help with specimen collections in Carrie Bow Cay. We thank Dr. Noah Palm of Yale University for use of his laboratory’s MiSeq, and to the Yale Center for Genome Analysis for their careful attention to the 2bRAD and RNA sequencing. This study was supported by a Fellowship of Graduate Student Travel (Society for Integrative and Comparative Biology) and Dr. David F. Ludwig Memorial Student Travel Scholarship (Association for Environmental Health and Sciences Foundation) awarded to J.B.K. and by grants awarded to R.W.T. from the U.S. National Science Foundation (DEB-1622398, DEB-1623837, OCE-1756249).

---

**Figure S1.**
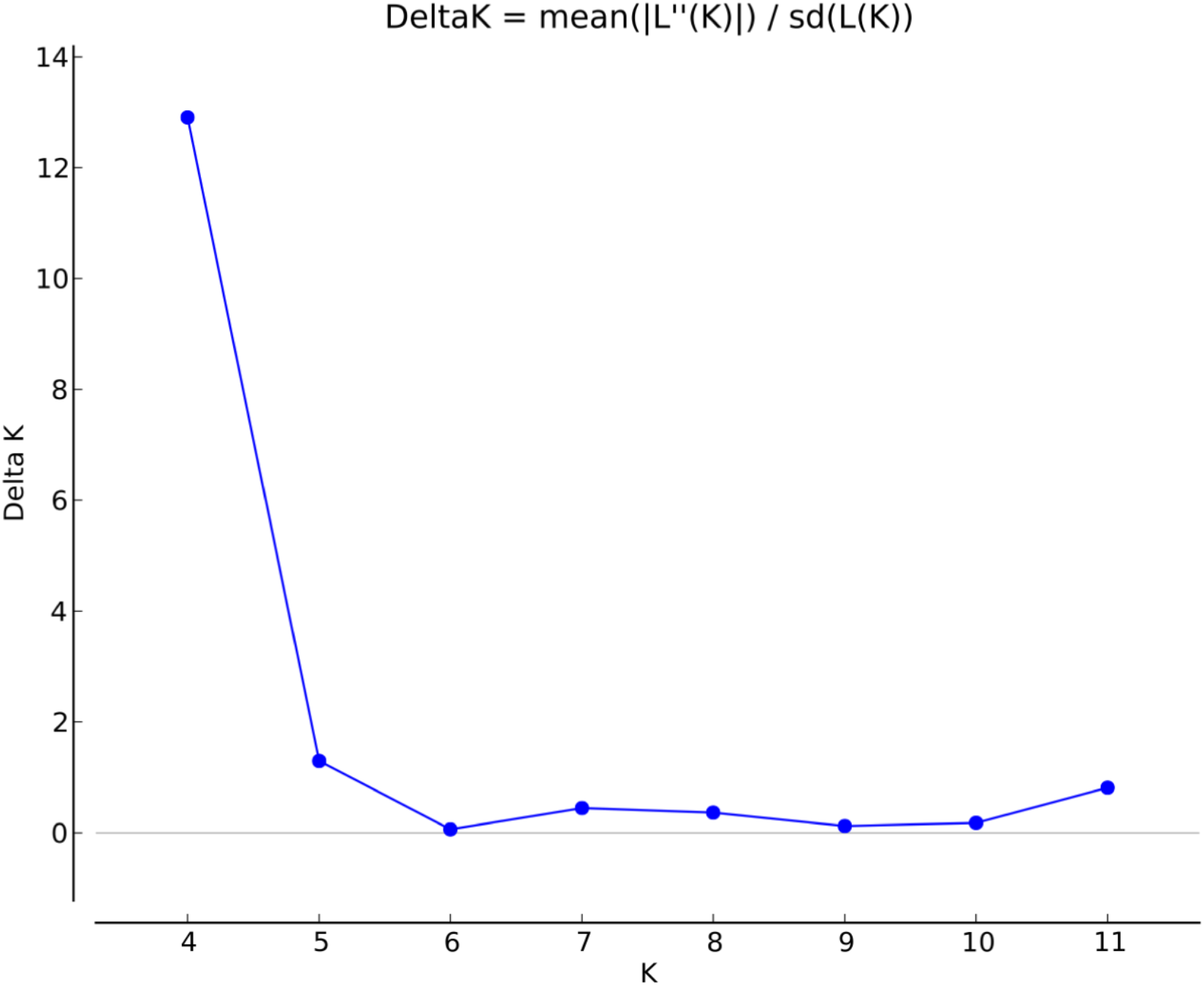
K=4 ancestral populations of *Ircinia* receive highest support via the Evanno method, followed by K=5. Estimation of delta K using Structure Harvester.

## References

1. Haney CH, Samuel BS, Bush J, Ausubel FM. 2015 Associations with rhizosphere bacteria can confer an adaptive advantage to plants. Nat. Plants 1, 15051. (doi:10.1038/nplants.2015.51)

2. Berendsen RL, Pieterse CMJ, Bakker PAHM. 2012 The rhizosphere microbiome and plant health. Trends Plant Sci. 17, 478–486. (doi:10.1016/j.tplants.2012.04.001)

3. Clemente JC, Ursell LK, Parfrey LW, Knight R. 2012 The Impact of the Gut Microbiota on Human Health: An Integrative View. Cell 148, 1258–1270. (doi:10.1016/j.cell.2012.01.035)

4. Freeman CJ, Thacker RW. 2011 Complex interactions between marine sponges and their symbiotic microbial communities. Limnol. Oceanogr. 56, 1577–1586. (doi:10.4319/lo.2011.56.5.1577)

5. Moran NA, Baumann P. 2000 Bacterial endosymbionts in animals. Curr. Opin. Microbiol. 3, 270–275. (doi:https://doi.org/10.1016/S1369-5274(00)00088-6)

6. Falkowski PG, Dubinsky Z, Muscatine L, Porter JW. 1984 Light and the Bioenergetics of a Symbiotic Coral. Bioscience 34, 705–709. (doi:10.2307/1309663)

7. Long SR. 1989 Rhizobium-legume nodulation: Life together in the underground. Cell 56, 203–214. (doi:https://doi.org/10.1016/0092-8674(89)90893-3)

8. Yeoman CJ, White BA. 2014 Gastrointestinal Tract Microbiota and Probiotics in Production Animals. Annu. Rev. Anim. Biosci. 2, 469–486. (doi:10.1146/annurev-animal-022513-114149)

9. Brown SP, Cornforth DM, Mideo N. 2012 Evolution of virulence in opportunistic pathogens: generalism, plasticity, and control. Trends Microbiol. 20, 336–342. (doi:10.1016/j.tim.2012.04.005)

10. Kiers ET, Rousseau RA, West SA, Denison RF. 2003 Host sanctions and the legume-rhizobium mutualism. Nature 425, 78–81. (doi:10.1038/nature01931)

11. Aderem A, Underhill DM. 1999 Mechanisms of Phagocytosis in Macrophages. Annu. Rev. Immunol. 17, 593–623. (doi:10.1146/annurev.immunol.17.1.593)

12. Nyholm S V, McFall-Ngai M. 2004 The winnowing: establishing the squid–vibrio symbiosis. Nat. Rev. Microbiol. 2, 632–642. (doi:10.1038/nrmicro957)

13. Wiens M, Korzhev M, Perovic-Ottstadt S, Luthringer B, Brandt D, Klein S, Muller WEG. 2007 Toll-like receptors are part of the innate immune defense system of sponges (demospongiae: Porifera). Mol. Biol. Evol. 24, 792–804. (doi:10.1093/molbev/msl208)

14. Sweet MJ, Hume DA. 1996 Endotoxin signal transduction in macrophages. J. Leukoc. Biol. 60, 8–26. (doi:10.1002/jlb.60.1.8)

15. Böhm M, Hentschel U, Friedrich A, Fieseler L, Steffen R, Gamulin V, Müller I, Müller W. 2001 Molecular response of the sponge Suberites domuncula to bacterial infection. Mar. Biol. 139, 1037–1045. (doi:10.1007/s002270100656)

16. Dunn CW, Leys SP, Haddock SHD. 2015 The hidden biology of sponges and ctenophores. Trends Ecol. Evol. 30, 282–291. (doi:10.1016/j.tree.2015.03.003)

17. Brunton FR, Dixon OA. 1994 Siliceous sponge-microbe biotic associations and their recurrence through the Phanerozoic as reef mound constructors. Palaios 9, 370–387. (doi:10.2307/3515056)

18. Vacelet J, Donadey C. 1977 Vacelet J, Donadey C.. Electron microscope study of the association between some sponges and bacteria. J Exp Mar Biol Ecol 30:301–314. (doi:10.1016/0022-0981(77)90038-7)

19. Wilkinson CR. 1978 Microbial associations in sponges. II. Numerical analysis of sponge and water bacterial populations. Mar. Biol. 49, 169–176. (doi:10.1007/BF00387116)

20. Hentschel U, Usher KM, Taylor MW. 2006 Marine sponges as microbial fermenters. FEMS Microbiol. Ecol. 55, 167–177. (doi:10.1111/j.1574-6941.2005.00046.x)

21. Thomas T et al. 2016 Diversity, structure and convergent evolution of the global sponge microbiome. Nat. Commun. 7, 11870.

22. Thomas TRA, Kavlekar DP, LokaBharathi PA. 2010 Marine drugs from sponge-microbe association--a review. Mar. Drugs 8, 1417–1468. (doi:10.3390/md8041417)

23. Tian R-M, Wang Y, Bougouffa S, Gao Z-M, Cai L, Bajic V, Qian P-Y. 2014 Genomic analysis reveals versatile heterotrophic capacity of a potentially symbiotic sulfur-oxidizing bacterium in sponge. Environ. Microbiol. 16, 3548–3561. (doi:10.1111/1462-2920.12586)

24. Archer SK, Stevens JL, Rossi RE, Matterson KO, Layman CA. 2017 Abiotic conditions drive significant variability in nutrient processing by a common Caribbean sponge, Ircinia felix. Limnol. Oceanogr. 62, 1783–1793. (doi:10.1002/lno.10533)

25. Slaby BM, Hackl T, Horn H, Bayer K, Hentschel U. 2017 Metagenomic binning of a marine sponge microbiome reveals unity in defense but metabolic specialization. Nat. Publ. Gr. 11, 2465–2478. (doi:10.1038/ismej.2017.101)

26. Rix L, Ribes M, Coma R, Jahn MT, de Goeij JM, van Oevelen D, Escrig S, Meibom A, Hentschel U. 2020 Heterotrophy in the earliest gut: a single-cell view of heterotrophic carbon and nitrogen assimilation in sponge-microbe symbioses. ISME J. (doi:10.1038/s41396-020-0706-3)

27. Achlatis M, Pernice M, Green K, Guagliardo P, Kilburn MR, Hoegh-Guldberg O, Dove S. 2018 Single-cell measurement of ammonium and bicarbonate uptake within a photosymbiotic bioeroding sponge. ISME J. 12, 1308–1318. (doi:10.1038/s41396-017-0044-2)

28. Erwin PM, Thacker RW. 2008 Phototrophic nutrition and symbiont diversity of two Caribbean sponge – cyanobacteria symbioses. (doi:10.3354/meps07464)

29. Wilson MC et al. 2014 An environmental bacterial taxon with a large and distinct metabolic repertoire. Nature 506, 58–62. (doi:10.1038/nature12959)

30. Lackner G, Peters EE, Helfrich EJN, Piel J. 2017 Insights into the lifestyle of uncultured bacterial natural product factories associated with marine sponges. Proc. Natl. Acad. Sci. 114, E347 LP–E356. (doi:10.1073/pnas.1616234114)

31. Moitinho-Silva L et al. 2017 The sponge microbiome project. Gigascience 6, 1–7. (doi:10.1093/gigascience/gix077)

32. Freeman CJ, Easson CG, Matterson KO, Thacker RW, Baker DM, Paul VJ. 2020 Microbial symbionts and ecological divergence of Caribbean sponges: A new perspective on an ancient association. ISME J. (doi:10.1038/s41396-020-0625-3)

33. Erwin PM, Pita L, López-Legentil S, Turon X. 2012 Stability of sponge-associated bacteria over large seasonal shifts in temperature and irradiance. Appl. Environ. Microbiol. 78, 7358–7368. (doi:10.1128/AEM.02035-12)

34. Marino C, Pawlik J, Lopez-Legentil S, Erwin P. 2017 Latitudinal variation in the microbiome of the sponge Ircinia campana correlates with host haplotype but not anti-predatory chemical defense. Mar. Ecol. Prog. Ser. 565, 53–66. (doi:10.3354/meps12015)

35. Griffiths S, Antwis R, Lenzi L, Lucaci A, Behringer DC, Butler M, Preziosi R. 2019 Host genetics and geography influence microbiome composition in the sponge Ircinia campana. J. Anim. Ecol. 88, 1684–1695. (doi:10.1111/1365-2656.13065)

36. Van Soest RWM et al. 2019 World Porifera Database. (doi:10.14284/359)

37. Pita L, Turon X, López-Legentil S, Erwin PM. 2013 Host rules: spatial stability of bacterial communities associated with marine sponges (Ircinia spp.) in the Western Mediterranean Sea. FEMS Microbiol. Ecol. 86, 268–276. (doi:10.1111/1574-6941.12159)

38. Wilkinson C, Cheshire A. 1990 Comparisons of sponge populations across the Barrier Reefs of Australia and Belize: evidence for higher productivity in the Caribbean. Mar. Ecol. Prog. Ser. 67, 285–294. (doi:10.3354/meps067285)

39. Weisz JB, Hentschel U, Lindquist N, Martens CS. 2007 Linking abundance and diversity of sponge-associated microbial communities to metabolic differences in host sponges. Mar. Biol. 152, 475–483. (doi:10.1007/s00227-007-0708-y)

40. Schloss PD et al. 2009 Introducing mothur: open-source, platform-independent, community-supported software for describing and comparing microbial communities. Appl. Environ. Microbiol. 75, 7537–7541. (doi:10.1128/AEM.01541-09)

41. Anderson MJ. 2005 PERMANOVA Permutational multivariate analysis of variance. Austral Ecol. (doi:10.1139/cjfas-58-3-626)

42. Jackson AL, Parnell AC, Inger R, Bearhop S. 2011 Comparing isotopic niche widths among and within communities: SIBER – Stable Isotope Bayesian Ellipses in R. J. Anim. Ecol., 595–602.

43. Bolger AM, Lohse M, Usadel B. 2014 Genome analysis Trimmomatic: a flexible trimmer for Illumina sequence data. Bioinformatics 30, 2114–2120. (doi:10.1093/bioinformatics/btu170)

44. Quast C, Pruesse E, Yilmaz P, Gerken J, Schweer T, Yarza P, Peplies J, Glöckner FO. 2013 The SILVA ribosomal RNA gene database project: Improved data processing and web-based tools. Nucleic Acids Res. 41, D590–D596. (doi:10.1093/nar/gks1219)

45. Yamada K, Fukuda W, Kondo Y, Miyoshi Y, Atomi H, Imanaka T. 2011 Constrictibacter antarcticus gen. nov., sp. nov., a cryptoendolithic micro-organism from Antarctic white rock. Int. J. Syst. Evol. Microbiol. 61, 1973–1980. (doi:10.1099/ijs.0.026625-0)

46. Flanagan SP, Jones AG. 2017 Constraints on the FST–Heterozygosity Outlier Approach. J. Hered. 108, 561–573. (doi:10.1093/jhered/esx048)

47. Foll M, Gaggiotti O. 2008 A Genome-Scan Method to Identify Selected Loci Appropriate for Both Dominant and Codominant Markers: A Bayesian Perspective. Genetics 180, 977LP–993. (doi:10.1534/genetics.108.092221)

48. Wood DE, Salzberg SL. 2014 Kraken: ultrafast metagenomic sequence classification using exact alignments. Genome Biol. 15, R46. (doi:10.1186/gb-2014-15-3-r46)

49. Schmieder R, Edwards R. 2011 Fast identification and removal of sequence contamination from genomic and metagenomic datasets. PLoS One 6, e17288. (doi:10.1371/journal.pone.0017288)

50. Martin M. 2011 CUTADAPT removes adapter sequences from high-throughput sequencing reads. EMBnet.journal 17. (doi:10.14806/ej.17.1.200)

51. Catchen J, Hohenlohe PA, Bassham S, Amores A, Cresko WA. 2013 Stacks: an analysis tool set for population genomics. Mol. Ecol. 22, 3124–3140. (doi:10.1111/mec.12354)

52. Grabherr MG et al. 2011 Full-length transcriptome assembly from RNA-Seq data without a reference genome. Nat. Biotechnol. (doi:10.1038/nbt.1883)

53. Scott C. 2016 dammit: an open and accessible de novo transcriptome annotator. in prep.

54. Leaché AD, Fujita MK, Minin VN, Bouckaert RR. 2014 Species delimitation using genome-wide SNP Data. Syst. Biol. 63, 534–542. (doi:10.1093/sysbio/syu018)

55. Kass RE, Raftery AE. 1995 Bayes Factors. J. Am. Stat. Assoc. 90, 773–795. (doi:10.1080/01621459.1995.10476572)

56. Evanno G, Regnaut S, Goudet J. 2005 Detecting the number of clusters of individuals using the software STRUCTURE: a simulation study. Mol. Ecol. 14, 2611–2620. (doi:10.1111/j.1365-294X.2005.02553.x)

57. Liu L, Yu L, Kubatko L, Pearl DK, Edwards S V. 2009 Coalescent methods for estimating phylogenetic trees. Mol. Phylogenet. Evol. 53, 320–328. (doi:https://doi.org/10.1016/j.ympev.2009.05.033)

58. Schmitt S, Weisz JB, Lindquist N, Hentschel U. 2007 Vertical transmission of a phylogenetically complex microbial consortium in the viviparous sponge Ircinia felix. Appl. Environ. Microbiol. 73, 2067–2078. (doi:10.1128/AEM.01944-06)

59. Lee OO, Chui PY, Wong YH, Pawlik JR, Qian P-Y. 2009 Evidence for vertical transmission of bacterial symbionts from adult to embryo in the Caribbean sponge Svenzea zeai. Appl. Environ. Microbiol. 75, 6147–6156. (doi:10.1128/AEM.00023-09)

60. Sharp KH, Eam B, Faulkner DJ, Haygood MG. 2007 Vertical Transmission of Diverse Microbes in the Tropical Sponge *Corticium* sp. Appl. Environ. Microbiol. 73, 622 LP–629. (doi:10.1128/AEM.01493-06)

61. Enticknap JJ, Kelly M, Peraud O, Hill RT. 2006 Characterization of a culturable alphaproteobacterial symbiont common to many marine sponges and evidence for vertical transmission via sponge larvae. Appl. Environ. Microbiol. 72, 3724–3732. (doi:10.1128/AEM.72.5.3724-3732.2006)

62. Erwin PM, Thacker RW. 2007 Incidence and identity of photosynthetic symbionts in Caribbean coral reef sponge assemblages. J. Mar. Biol. Assoc. United Kingdom 87, 1683–1692. (doi:10.1017/S0025315407058213)

63. Glasl B, Smith CE, Bourne DG, Webster NS. 2018 Exploring the diversity-stability paradigm using sponge microbial communities. Sci. Rep. 8, 8425. (doi:10.1038/s41598-018-26641-9)

64. Ferrer-Admetlla A et al. 2008 Balancing Selection Is the Main Force Shaping the Evolution of Innate Immunity Genes. J. Immunol. 181, 1315 LP–1322. (doi:10.4049/jimmunol.181.2.1315)

65. Leiva C, Taboada S, Kenny NJ, Combosch DJ, Giribet G, Jombart T, Riesgo A. 2019 Population substructure and signals of divergent adaptive selection despite admixture in the sponge Dendrilla antarctica from shallow waters surrounding the Antarctic Peninsula. Mol. Ecol. 28, 3151–3170.

66. Orton RJ, Sturm OE, Vyshemirsky V, Calder M, Gilbert DR, Kolch W. 2005 Computational modelling of the receptor-tyrosine-kinase-activated MAPK pathway. Biochem. J. 392, 249–261. (doi:10.1042/BJ20050908)

67. Vos MD, Ellis CA, Bell A, Birrer MJ, Clark GJ. 2000 Ras Uses the Novel Tumor Suppressor RASSF1 as an Effector to Mediate Apoptosis. J. Biol. Chem. 275, 35669–35672. (doi:10.1074/jbc.C000463200)

68. Agathanggelou A, Cooper WN, Latif F. 2005 Role of the Ras-Association Domain Family 1 Tumor Suppressor Gene in Human Cancers. Cancer Res. 65, 3497 LP–3508. (doi:10.1158/0008-5472.CAN-04-4088)

69. Donninger H, Vos MD, Clark GJ. 2007 The RASSF1A tumor suppressor. J. Cell Sci. 120, 3163 LP–3172. (doi:10.1242/jcs.010389)

70. Stark MS et al. 2011 Frequent somatic mutations in MAP3K5 and MAP3K9 in metastatic melanoma identified by exome sequencing. Nat. Genet. 44, 165–169. (doi:10.1038/ng.1041)

71. Coutts AS, MacKenzie E, Griffith E, Black DM. 2003 TES is a novel focal adhesion protein with a role in cell spreading. J. Cell Sci. 116, 897–906. (doi:10.1242/jcs.00278)

72. Manton KJ, Douglas ML, Netzel-Arnett S, Fitzpatrick DR, Nicol DL, Boyd AW, Clements JA, Antalis TM. 2005 Hypermethylation of the 5’ CpG island of the gene encoding the serine protease Testisin promotes its loss in testicular tumorigenesis. Br. J. Cancer 92, 760–769. (doi:10.1038/sj.bjc.6602373)

73. Li X et al. 2014 Histone demethylase KDM5B is a key regulator of genome stability. Proc. Natl. Acad. Sci. 111, 7096 LP–7101. (doi:10.1073/pnas.1324036111)

74. Brachner A et al. 2012 The endonuclease Ankle1 requires its LEM and GIY-YIG motifs for DNA cleavage in vivo. J. Cell Sci. 125, 1048–1057. (doi:10.1242/jcs.098392)

75. Dunin-Horkawicz S, Feder M, Bujnicki JM. 2006 Phylogenomic analysis of the GIY-YIG nuclease superfamily. BMC Genomics 7, 98. (doi:10.1186/1471-2164-7-98)

76. Suzuki T, Yasui M, Honma M. 2016 Mutator Phenotype and DNA Double-Strand Break Repair in BLM Helicase-Deficient Human Cells. Mol. Cell. Biol. 36, 2877–2889. (doi:10.1128/MCB.00443-16)

77. Kalinovskaya NI et al. 1995 Surfactin-like structures of five cyclic depsipeptides from the marine isolate of Bacillus pumilus. Russ. Chem. Bull. 44, 951–955. (doi:10.1007/BF00696935)

78. Prokof’eva NG, Kalinovskaya NI, Luk’yanov PA, Shentsova EB, Kuznetsova TA. 1999 The membranotropic activity of cyclic acyldepsipeptides from bacterium Bacillus pumilus, associated with the marine sponge Ircinia sp. Toxicon 37, 801–813. (doi:10.1016/s0041-0101(98)00219-0)

79. Hardoim CCP, Costa R. 2014 Microbial communities and bioactive compounds in marine sponges of the family Irciniidae-a review. Mar. Drugs 12, 5089–5122. (doi:10.3390/md12105089)

80. Jahn M et al. 2019 A Phage Protein Aids Bacterial Symbionts in Eukaryote Immune Evasion. Cell Host Microbe 26. (doi:10.1016/j.chom.2019.08.019)

81. González-Morales N, Holenka TK, Schöck F. 2017 Filamin actin-binding and titin-binding fulfill distinct functions in Z-disc cohesion. PLOS Genet. 13, e1006880.

82. Leber Y, Ruparelia AA, Kirfel G, van der Ven PFM, Hoffmann B, Merkel R, Bryson-Richardson RJ, Fürst DO. 2016 Filamin C is a highly dynamic protein associated with fast repair of myofibrillar microdamage. Hum. Mol. Genet. 25, 2776–2788. (doi:10.1093/hmg/ddw135)

83. Mao Z, Nakamura F. 2020 Structure and Function of Filamin C in the Muscle Z-Disc. Int. J. Mol. Sci. 21. (doi:10.3390/ijms21082696)

84. Leys SP, Yahel G, Reidenbach MA, Tunnicliffe V, Shavit U, Reiswig HM. 2011 The sponge pump: the role of current induced flow in the design of the sponge body plan. PLoS One 6, e27787–e27787. (doi:10.1371/journal.pone.0027787)

85. Hoffmann F, Røy H, Bayer K, Hentschel U, Pfannkuchen M, Brümmer F, de Beer D. 2008 Oxygen dynamics and transport in the Mediterranean sponge Aplysina aerophoba. Mar. Biol. 153, 1257–1264. (doi:10.1007/s00227-008-0905-3)

86. Guannel G, Arkema K, Ruggiero P, Verutes G. 2016 The Power of Three: Coral Reefs, Seagrasses and Mangroves Protect Coastal Regions and Increase Their Resilience. PLoS One 11, e0158094.

87. Müller WEG, Müller IM. 2003 Origin of the Metazoan Immune System: Identification of the Molecules and Their Functions in Sponges1. Integr. Comp. Biol. 43, 281–292. (doi:10.1093/icb/43.2.281)

88. Wang S, Meyer E, Mckay JK, Matz M V. 2012 2b-RAD: A simple and flexible method for genome-wide genotyping. Nat. Methods 9, 808–810. (doi:10.1038/nmeth.2023)

89. Chen S, Zhou Y, Chen Y, Gu J. 2018 Fastp: an ultra-fast all-in-one FASTQ preprocessor. Bioinformatics 34, i884–i890. (doi:10.1093/bioinformatics/bty560)

90. Rognes T, Flouri T, Nichols B, Quince C, Mahé F. 2016 VSEARCH: a versatile open source tool for metagenomics. PeerJ 4, e2584–e2584. (doi:10.7717/peerj.2584)

91. Oksanen AJ et al. 2016 Vegan: community ecology package. https://github.com/vegandevs/vegan (doi:10.4135/9781412971874.n145)

92. Schmieder R, Edwards R. 2011 Quality control and preprocessing of metagenomic datasets. Bioinformatics 27, 863–864. (doi:10.1093/bioinformatics/btr026)

93. Paris JR, Stevens JR, Catchen JM. 2017 Lost in parameter space: a road map for stacks. Methods Ecol. Evol. 8, 1360–1373. (doi:10.1111/2041-210X.12775)

94. Beaumont MA, Nichols RA. 1996 Evaluating Loci for Use in the Genetic Analysis of Population Structure. Proc. R. Soc. London Ser. B 263, 1619–1626.

95. Bouckaert R, Heled J, Kühnert D, Vaughan T, Wu CH, Xie D, Suchard MA, Rambaut A, Drummond AJ. 2014 BEAST 2: a software platform for Bayesian evolutionary analysis. PLoS Comput. Biol. 10, e1003537. (doi:10.1371/journal.pcbi.1003537)

96. Bryant D, Bouckaert R, Felsenstein J, Rosenberg NA, Roychoudhury A. 2012 Inferring species trees directly from biallelic genetic markers: bypassing gene trees in a full coalescent analysis. Mol. Biol. Evol. 29, 1917–1932. (doi:10.1093/molbev/mss086)

97. Earl D, vonHoldt BM. 2011 STRUCTURE HARVESTER: a website and program for visualizing STRUCTURE output and implementing the Evanno method. Conserv. Genet. Resour. 4, 359–361.

